# NINJ1 mediates hepatic ischemia-reperfusion injury

**DOI:** 10.64898/2026.02.07.704585

**Authors:** J. Mossemann, B. Martins, Y. Zhao, F. Aguilar, D. Taskina, C.J. Hur, A. Volchuk, G. Ye, D.M. Ali, A. Mirzaesmaeili, I. Siddiqui, G. Goodarzi, P.J. Bilan, I.B. Stowe, N. Kayagaki, S. MacParland, S. Freeman, N.M. Goldenberg, B.E. Steinberg, B.A. Sayed

## Abstract

Hepatic ischemia-reperfusion injury (IRI) results from interrupted perfusion to the liver and contributes to acute liver dysfunction such as following liver transplantation. Lytic cell death pathways are major drivers of IRI and the subsequent inflammatory response. The transmembrane protein ninjurin-1 (NINJ1) was identified as the key executor of terminal plasma membrane rupture across multiple lytic cell death pathways implicated in hepatic IRI. We hypothesized that NINJ1-mediated lytic cell death drives IRI and that its therapeutic inhibition would mitigate liver IRI. Using human liver specimens, we found that NINJ1 is highly expressed in human liver tissue and that its activation correlates with early allograft dysfunction in patients undergoing liver transplantation. Utilizing a segmental hepatic IRI model in mice and rats, *Ninj1* genetic deletion or pharmacologic inhibition diminished acute liver injury. Mice with hepatocyte- or macrophage-specific *Ninj1* knockout both had reduced hepatocellular injury following IRI, suggesting that NINJ1 within both populations contributes to the resulting liver injury. Mechanistically, we found that hepatocytes and Kupffer cells are highly susceptible to hypoxia-induced NINJ1-mediated plasma membrane rupture, which can be pharmacologically prevented. These data position NINJ1 as a potential new therapeutic target to limit hepatic IRI, with important implications for organ preservation during liver transplantation.

## Introduction

Hepatic ischemia reperfusion injury (IRI) is a two-hit phenomenon caused by temporarily interrupting liver perfusion (ischemia), followed by the re-establishment of normal blood flow (reperfusion). IRI occurs in a variety of settings, the most common being during liver transplantation and hemodynamic shock. Ischemia disrupts cellular homeostasis to activate both necrotic and regulated forms of cell death (1, 2). Many of these dying cells lose structural integrity, thereby releasing cytoplasmic damage-associated molecular patterns (DAMPs), reactive oxygen species and other inflammatory mediators into the local tissue environment. When blood flow is restored, the cytotoxic molecules released from dying cells spread throughout the tissue and into the systemic circulation, amplifying inflammation, secondary cellular stress and ultimately further propagating lytic cell death (3). Together, the local and systemic sequalae of hepatic IRI manifest as ischemia-reperfusion syndrome, a transient liver insufficiency and systemic sterile inflammatory response that drive acute patient morbidity and mortality (3, 4). Severe acute IRI can also lead to long-term debilitating complications including liver fibrosis and, in the setting of transplant, predisposition to rejection (5).

Regulated lytic cell death pathways are important contributors to human liver disease (6–8), a finding recapitulated in animal models (9, 10). The specific lytic cell death pathways activated in liver IRI include secondary necrosis, the breakdown of apoptotic bodies that are not removed by phagocytes at the site of injury (11), and pyroptosis, a pro-inflammatory death pathway that depends on the activation of pore-forming gasdermin proteins (12). Necroptosis and ferroptosis are other forms of lytic cell death implicated in hepatic IRI, especially in patients with co-morbidities such as hepatic steatosis, advanced age or iron overload (13–16). In each of secondary necrosis, pyroptosis and ferroptosis, the terminal event of plasma membrane rupture is dependent on the protein ninjurin-1 (NINJ1) (17–19). NINJ1 is a 16-kDa transmembrane protein with particularly high expression in the liver and innate immune cells (20, 21). As NINJ1-expressing cells undergo cell death via these pathways, NINJ1 clusters into high-order oligomers within the plasma membrane to destabilize and disrupt the plasma membrane (22–24).

Early evidence with an anti-NINJ1 neutralizing antibody, termed clone D1, indicates that preventing NINJ1 clustering mitigates liver injury in multiple mouse models of acute injury, including IRI (25). As compared to isotype control, clone D1 administration 4 hours prior to hepatic IRI reduced circulating markers of hepatocellular injury and liver cell death (25). These findings are consistent with the known role of plasma membrane rupture in the multiple cell death pathways that propagate liver damage in IRI. Similarly, the cytoprotective amino acid glycine and small molecule muscimol preserve cellular integrity by inhibiting NINJ1 activation (26, 27), positioning NINJ1 as a viable therapeutic target. We therefore hypothesized that NINJ1 activation occurs in human IRI and that its genetic deletion or pharmacologic inhibition would protect the liver from injury in animal models. Here, we demonstrate that NINJ1 is expressed in human liver and that human liver allografts with severe early allograft dysfunction post-transplant have increased activation of NINJ1. In both mouse and rat *Ninj1* knockout models, the absence of NINJ1 diminishes injury from IRI, substantiating its role in driving hepatocellular damage. Compartmental NINJ1 deletion in hepatocytes or macrophages confer protection against acute IRI, establishing a role for NINJ1 in both cell types in driving IRI, a finding which we substantiate in an *in vitro* hypoxia-reoxygenation model. Together, our findings demonstrate a crucial role of NINJ1 in mediating acute hepatic IRI and indicate that targeting NINJ1 as a therapeutic target may dampen the severity of hepatic IRI.

## Results

### NINJ1 activation correlates with severe early allograft dysfunction following human liver transplantation

Genome Wide Association Studies have identified a human *NINJ1* single nucleotide polymorphism (rs7018885) which is associated with decreased liver transaminase levels (25); however, a role in human liver transplantation has not been previously explored. Using a cell type-annotated single cell RNA-sequencing dataset from human hepatic allografts utilized for transplantation (15), we examined NINJ1 transcript expression and distribution across immune and parenchymal clusters on UMAP embeddings (**Figure 1A-D**). *NINJ1* expression most enriched within myeloid populations, with prominent expression in Kupffer cells, as well as in parenchymal hepatocyte populations (**Figure 1A-B**). We next interrogated non-dissociative spatial transcriptomic data from intact liver sections, which confirmed cellular localization *in situ* but did not reveal a clear lobular zonation pattern for *NINJ1* expression (**Figure 1C-D**). We corroborated these spatial transcriptomics data using RNAscope *in situ* hybridization to examine the expression of *NINJ1* in healthy human liver samples. *NINJ1* mRNA was expressed across cell types and was particularly prominent in the parenchyma as compared to portal tracts (**Figure 1E**). *NINJ1* expression strongly co-localized with *CD68*, a macrophage and Kupffer cell marker, which were well visualized in the sinusoidal spaces (**Figure 1E**). *NINJ1* co-localized to a lesser extent with albumin (*ALB*)-expressing hepatocytes (**Figure 1E**).

**Figure 1.**
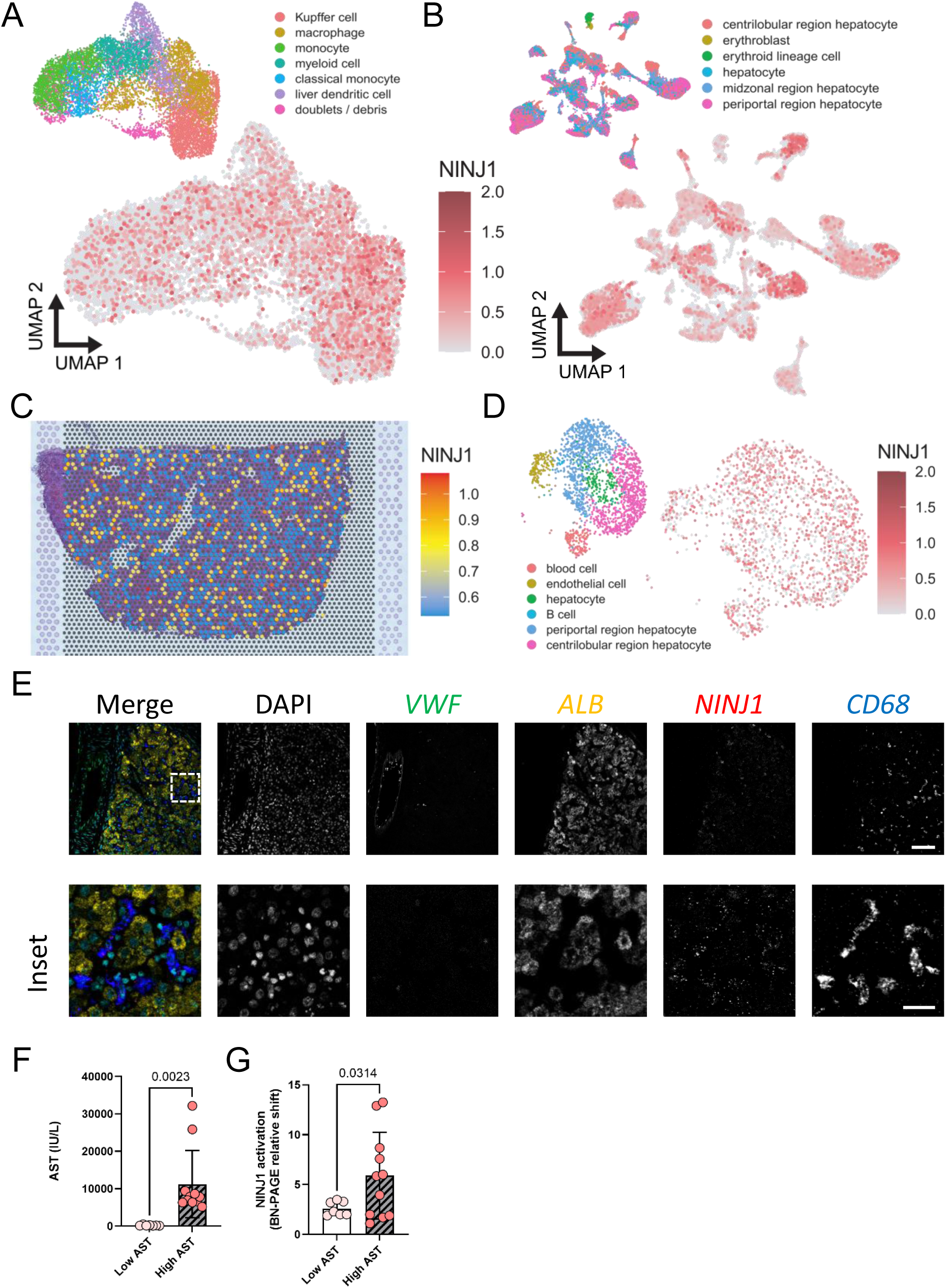
NINJ1 is expressed in the human liver and oligomerizes in ischemia-reperfusion injury. (**A-B**) Annotated UMAPs of single-cell RNA-sequencing atlas of non-diseased human livers (65) show NINJ1 transcripts across myeloid populations, with prominent expression in Kupffer cells, as well as in parenchymal hepatocytes. (**C-D**) Non-dissociative Visium spatial transcriptomics of an intact non-diseased human liver section from the same atlas (Andrews *et al.*) reveals capture spots over tissue confirming in situ expression of NINJ1, UMAP of Visium spots depict NINJ1 expression across different non- and parenchymal cells. (**E**) *In situ* hybridization using RNAscope of human liver shows co-expression of *NINJ1* (red) with markers of hepatocytes (*ALB*; yellow) and macrophages (*CD68*; blue). Endothelial cells were identified by expression of *VWF* (green). Nuclei stained with DAPI (cyan in merged image). Scale bar 100 microns (top row) and 30 microns for the inset (bottom row). (**F-G**) Patients following liver transplant were dichotomized into those with limited graft injury (aspartate aminotransferase, AST <500) and severe graft injury (AST > 5000). The extent of NINJ1 activation was assayed using blue native-PAGE on liver graft tissue. *P*-values were determined by T test.

Given the expression of NINJ1 in both Kupffer cell and hepatocyte populations, we next sought to specifically determine whether NINJ1 activation is linked with ischemia-reperfusion injury during liver transplantation. We examined transplanted human hepatic allografts (**Supplemental Table 1**) and characterized post-transplant graft injury based on serum markers of liver damage in the recipients that meet criteria for early allograft dysfunction based on serum aspartate aminotransferase levels (28).

Cryopreserved samples of human liver allografts post-reperfusion (prior to abdominal closure during primary transplant) were analyzed for NINJ1 activation by blue native-PAGE (**Figure 1G** and **Supplemental Figure 1**). With this approach, which maintains native protein complexes, activated and polymerized NINJ1 shifts to a series of high molecular weight bands (17, 26). We compared samples from patients with severe early allograft dysfunction (aspartate aminotransferase, AST >5000 U/L within 2 days post-transplant) versus those with limited evidence of IRI (AST <500 U/L within 2 days post-transplant) (**Figure 1F**). Post-reperfusion, there was minimal increase in NINJ1 oligomerization in the low injury group, but patients with severe early allograft dysfunction demonstrated significantly increased NINJ1 oligomerization evinced by aggregation into higher molecular weight species as compared to the minimal injury group (**Figure 1G** and **Supplemental Figure 1**). These data are consistent with enhanced NINJ1 clustering and activation in injured human liver allografts following transplantation.

### Hepatic ischemia-reperfusion injury activates NINJ1 in a mouse model

Having established that *NINJ1* is expressed in the human liver and that its activation correlates with heightened early allograft dysfunction following transplant, we turned to a mouse model to further delineate the mechanistic role of NINJ1 in IRI. In our model, mice were subjected to 70% segmental liver ischemia for 1 hour followed by 6 hours of reperfusion and compared to a sham laparotomy. In sham-operated animals, NINJ1 colocalized with both hepatocytes and Kupffer cells with NINJ1 expression enhancing in the hepatocyte compartment during IRI (**Figure 2A-B**). Blue native-PAGE of liver lysates from sham-treated mice showed that NINJ1 migrates as distinct dimers, tetramers and low-order oligomers (**Figure 2C**), consistent with the proposed resting oligomer state of NINJ1 (30). In contrast, IRI induced oligomerization of NINJ1 into higher molecular weight complexes, while diminishing the abundance of NINJ1 dimers (**Figure 2C**). Together, these data demonstrate that NINJ1 is expressed in hepatocyte and Kupffer cell populations and establish that NINJ1 is activated and has increased hepatocyte abundance during mouse IRI.

**Figure 2.**
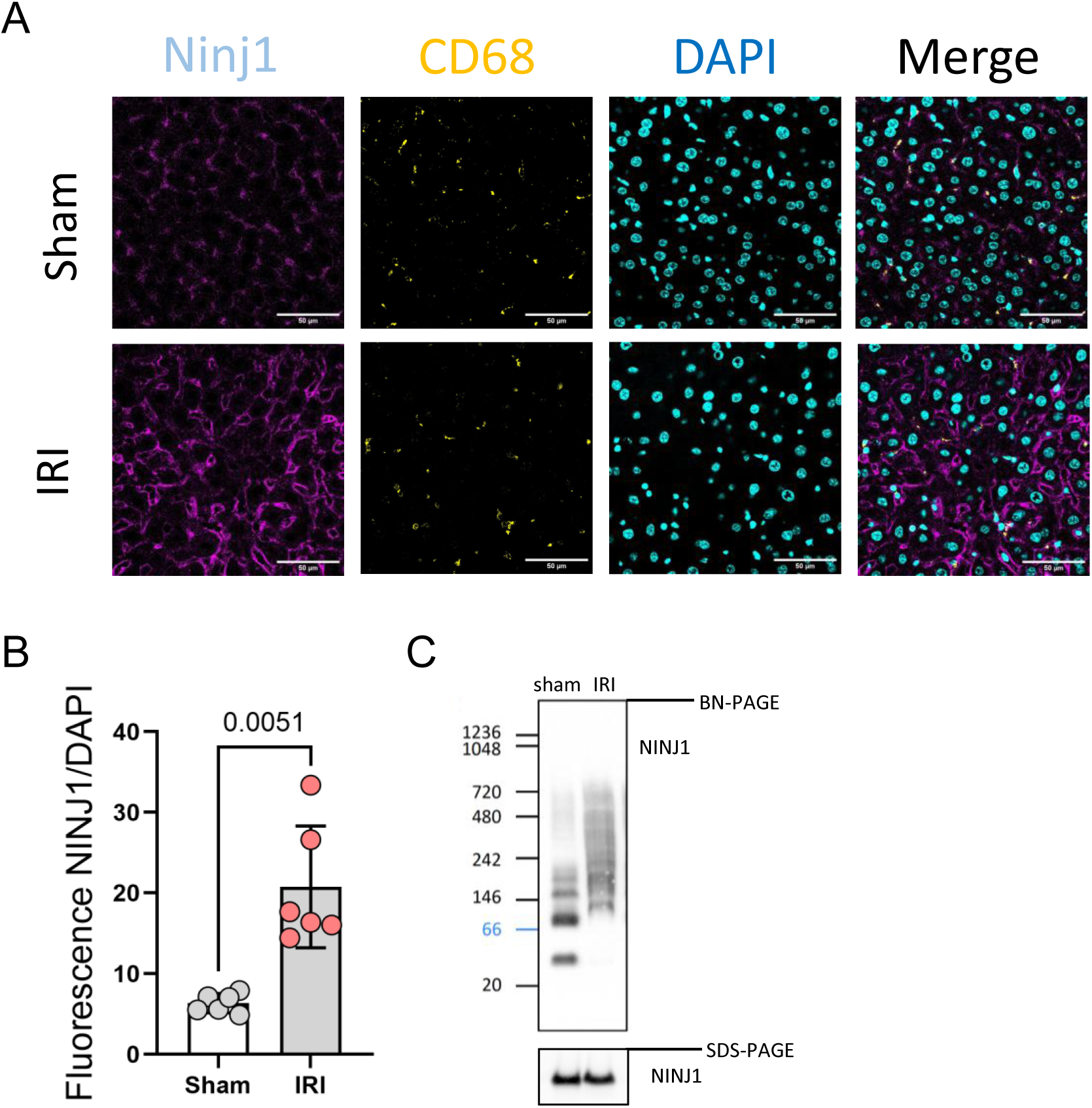
NINJ1 increases and oligomerizes following ischemia-reperfusion injury in a mouse model. Mixed-sex cohorts of wild-type mice underwent ischemia-reperfusion injury (IRI) consisting of 1 hour of 70% segmental warm ischemia followed by 6 hours of reperfusion prior to tissue collection. Sham laparotomy was used as a negative control. (**A**) Representative immunofluorescence of NINJ1 (magenta) in liver tissue from sham-operated or IRI liver. Kupffer cells were identified by CD68 (yellow). Scale bar 50 microns. (**B**) Quantification of total NINJ1 fluorescence in liver tissue from sham-operated (N = 6) and IRI (N = 6) animals as normalized to DAPI. *P*-value was determined by T test. (**C**) Representative blue native-PAGE for NINJ1 in liver lysates from sham-operated and IRI-treated animals. SDS-PAGE for NINJ1 provided as a loading control.

### *Ninj1* knockout limits mouse and rat hepatic ischemia-reperfusion injury

To determine if loss of NINJ1 offers protection from hepatic IRI, WT and whole-animal *Ninj1* knockout mice (*Ninj1^-/-^*) (17) (**Figure 3A**), were subjected to acute IRI. Serum lactate dehydrogenase (LDH; an indicator of cell rupture and non-specific tissue injury) and aminotransferases (AST and alanine aminotransferase, ALT; serum markers of hepatocyte injury) were used to evaluate the extent of liver injury (**Figure 3B-D**). Both WT and *Ninj1^-/-^* sham-operated mice did not show appreciable biochemical evidence of hepatocellular injury in their serum (**Figure 3B-D**). Hepatic IRI induced marked evidence of hepatocellular injury in WT mice, whereas *Ninj1^-/-^* mice had approximately 50% less biochemical evidence of liver injury following IRI compared with their WT counterparts (**Figure 3B-D**). Using the Suzuki scoring scale of histopathologic liver injury for IRI, the extent of liver injury was reduced in *Ninj1^-/-^*mice as compared to WT controls (31, 32) (**Figure 3F**), including a reduction of confluent necrosis (WT = 56.9 ± 4.9, N = 15; KO = 28.3 ± 9.6, N = 8; *P* = 0.0016; **Figure 3E,H**; **Supplemental Figure 2**). Finally, cell death was quantified using the TUNEL (terminal deoxynucleotidyl transferase dUTP nick end labeling) assay, which stains fragmented DNA to identify cells undergoing cell death (33) (**Figure 3G,I**; **Supplemental Figure 2**). Nearly one third of nuclei were TUNEL-positive in livers from WT mice injured by IRI, compared with 12 % TUNEL-positive nuclei in the *Ninj1*^-/-^ animals (**Figure 3G**). Notably, early inflammatory cell infiltration into the liver was not different between injured *Ninj1* WT and KO mice (**Supplemental Figure 3**). Minimal neutrophils, quantified as Ly6G-positive cells by immunohistochemistry, were observed in the livers of sham animals. In IRI mice, there was a comparable increase in neutrophil infiltration between genotypes (**Supplemental Figure 3A**). F4/80-positive macrophages resident in the liver did not change significantly in either genotype following IRI although there was a trend towards fewer macrophages (**Supplemental Figure 3B**) consistent with an observed early depletion of Kupffer cells in IRI (29).

**Figure 3.**
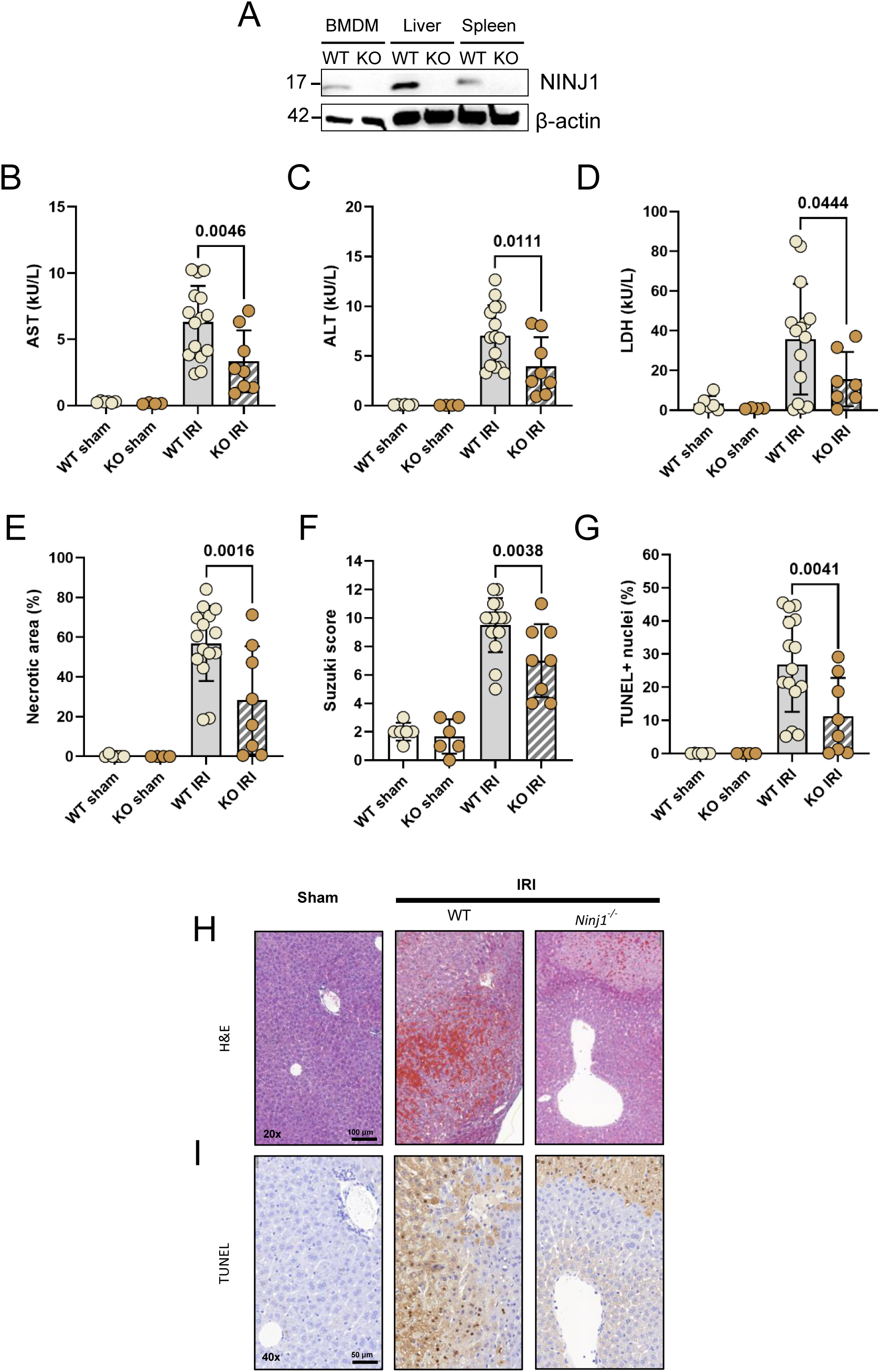
*Ninj1* knockout protects against liver ischemia-reperfusion injury in mice. Mixed-sex cohorts of *Ninj1* knockout mice or their littermate controls underwent ischemia-reperfusion injury (IRI) consisting of 1 hour of 70% segmental warm ischemia followed by 6 hours of reperfusion prior to tissue collection. Sham laparotomy was used as a negative control. Group sizes: Sham WT, N = 6; Sham KO, N = 4, IRI WT, N = 15; and IRI KO, N = 8. (**A**) SDS-PAGE showing NINJ1 expression in mouse bone marrow-derived macrophages, liver and spleen homogenates from wild-type animals compared with *Ninj1* knockout animals. (**B-D**) Hepatocyte-specific injury was assayed by serum AST and ALT, and general cellular cytotoxicity was measured by serum LDH. Note one missing data point in the KO IRI group for the LDH assay due to inadequate sample for analysis. Data points represent individual animals with the mean and standard error of the mean superimposed. *P*-values were determined by ANOVA with Tukey’s multiple comparison correction. (**E-I**) *Ninj1* knockout mice exposed to IRI had decreased liver tissue necrosis (**E**), Suzuki score (**F**) and cell death as assayed by TUNEL staining (**G**) compared to their littermate controls. Representative images of H&E (**H**) and TUNEL staining (**I**) are shown. Group sizes and statistical analysis the same as described for panels B-D, above. Data points represent individual animals with the mean and standard error of the mean superimposed. *P*-values were determined by ANOVA with Šidák’s multiple comparison correction.

We next evaluated whether the function of NINJ1 in IRI is conserved across species using *Ninj1^-/-^* KO rats (**Figure 4A**). *Ninj1^-/-^* rats had no evident basal phenotype, displaying normal home cage behaviour, and growth (**Supplemental Figure 4A**). To first determine if NINJ1 control of plasma membrane rupture is conserved in rats, bone-marrow derived macrophages (BMDMs) were cultured from *Ninj1^-/-^* and WT rats. Pyroptosis (LPS and nigericin) and secondary necrosis (ABT-199 and doxorubicin) were induced and plasma membrane rupture evaluated by LDH release into the supernatant. LDH release from *Ninj1^-/-^*BMDMs was significantly abrogated as compared to WT-derived cells, confirming NINJ1-dependent control of plasma membrane rupture in rat macrophages (**Supplemental Figure 4B**).

**Figure 4.**
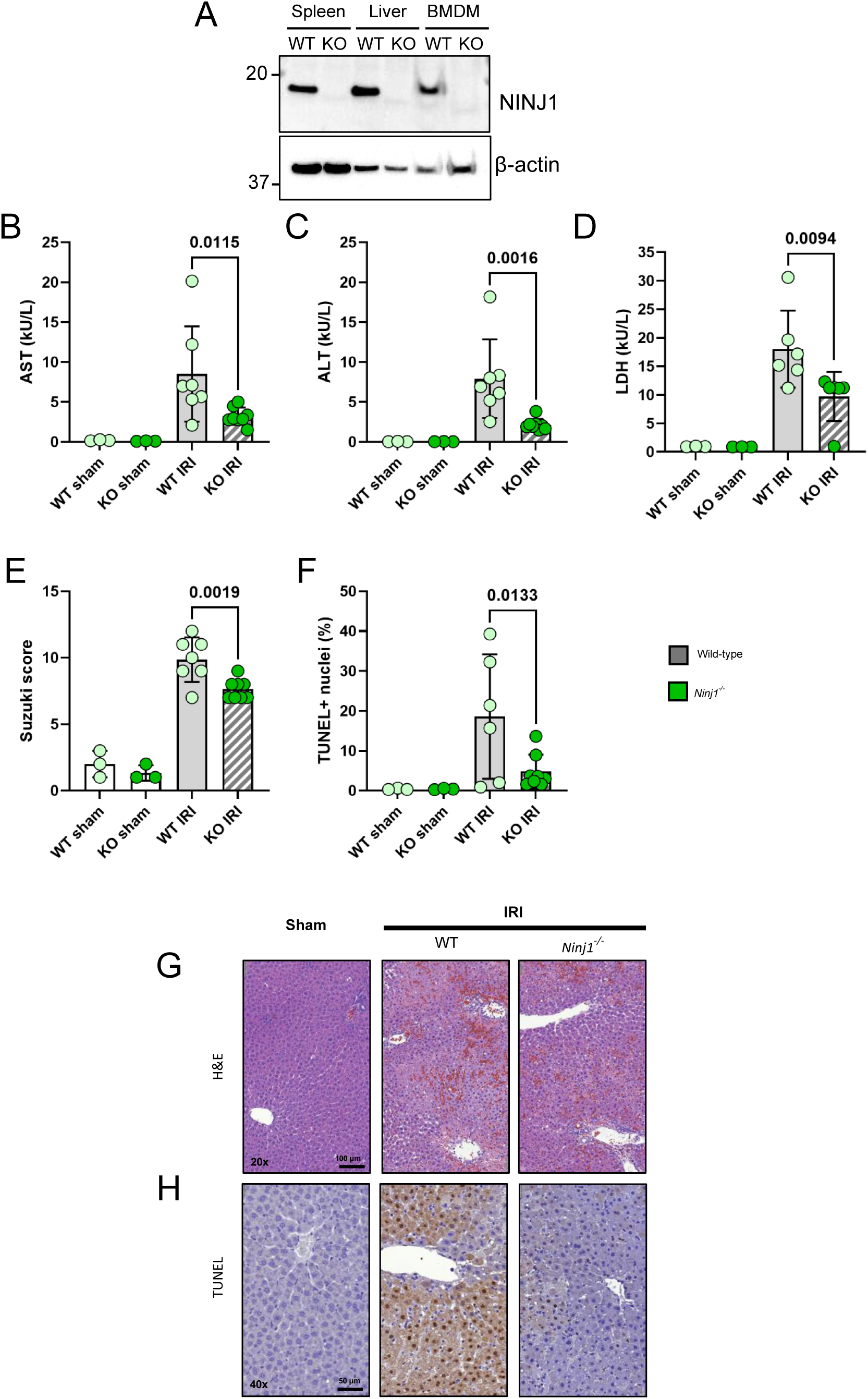
*Ninj1* knockout protects against liver ischemia-reperfusion injury in rats. Mixed-sex cohorts of *Ninj1* knockout rats or their littermate controls underwent ischemia-reperfusion injury (IRI) consisting of 1 hour of 70% segmental warm ischemia followed by 6 hours of reperfusion prior to tissue collection. Sham laparotomy was used as a negative control. Group sizes: Sham WT, N = 3; Sham KO, N = 3, IRI WT, N = 7; and IRI KO, N = 8. (**A**) SDS-PAGE demonstrating NINJ1 expression in rat bone marrow-derived macrophages, liver and spleen homogenates from wild-type animals compared with *Ninj1* knockout animals. (**B-D**) Hepatocyte-specific injury was assayed by serum AST and ALT, and general cellular cytotoxicity was measured by serum LDH. Data points represent individual animals with the mean and standard error of the mean superimposed. *P*-values were determined by ANOVA with Tukey’s multiple comparison correction. (**E-H**) *Ninj1* knockout rats exposed to IRI had decreased liver tissue necrosis and cell death compared to their littermate controls as shown by histopathology. (**E-H**) *Ninj1* knockout rats exposed to IRI had decreased Suzuki score (**E**) and cell death as assayed by TUNEL staining (**F**) compared to wild-type controls. Representative liver histology images of H&E (**G**) and TUNEL staining (**H**) are shown. Group sizes and statistical analysis the same as described for panels B-D, above. Data points represent individual animals with the mean and standard error of the mean superimposed. *P*-values were determined by ANOVA with Šidák’s multiple comparison correction.

The *Ninj1^-/-^* rats and their wild-type counterparts were subjected to a similar segmental model of ischemia (1 hour) and reperfusion (6 hours) as our mouse cohorts. *Ninj1^-/-^* rats had significant protection from hepatic IRI, as demonstrated by a reduction in AST, ALT and LDH compared to WT controls exposed to IRI (**Figure 4B-D**). Histopathologic analysis of the liver using the Suzuki score revealed decreased levels of necrosis, congestion and vacuolization in injured *Ninj1^-/-^*rats compared to WT (**Figure 4E,G**).

Additionally, cell death scored by TUNEL-positive nuclei, revealed a marked decrease in injured *Ninj1 ^-/-^*rats compared to WT (**Figure 4F,H**). These data in both mouse and rat indicate that NINJ1 deletion protects the liver against IRI across species.

### Pharmacologic NINJ1 inhibition with glycine reduces NINJ1 activation and hepatic ischemia-reperfusion injury in a mouse model

Empiric evidence supports the role of the amino acid glycine as a cytoprotective agent in organ preservation during IRI and transplant (34–39). We previously demonstrated that glycine cytoprotection works at the level of NINJ1, inhibiting its oligomerization and thus activation within the plasma membrane (26). To determine whether glycine protection from hepatic IRI correlates with NINJ1 inhibition, WT C57BL/6 mice were administered glycine by intraperitoneal injection one hour prior to the induction of hepatic ischemia and directly following the beginning of reperfusion. An equal dose of valine, an amino acid without cytoprotective properties (26, 40), was administered to WT mice as a placebo control. Glycine treatment also significantly lowered serum levels of AST, ALT and LDH induced by IRI compared to control mice (**Figure 5A-C**) and reduced confluent necrosis and percent TUNEL-positive nuclei relative to valine-treated animas (**Figure 5D-F**). Thus, pharmacologic inhibition of NINJ1 activation protects against liver injury following ischemia and reperfusion.

**Figure 5.**
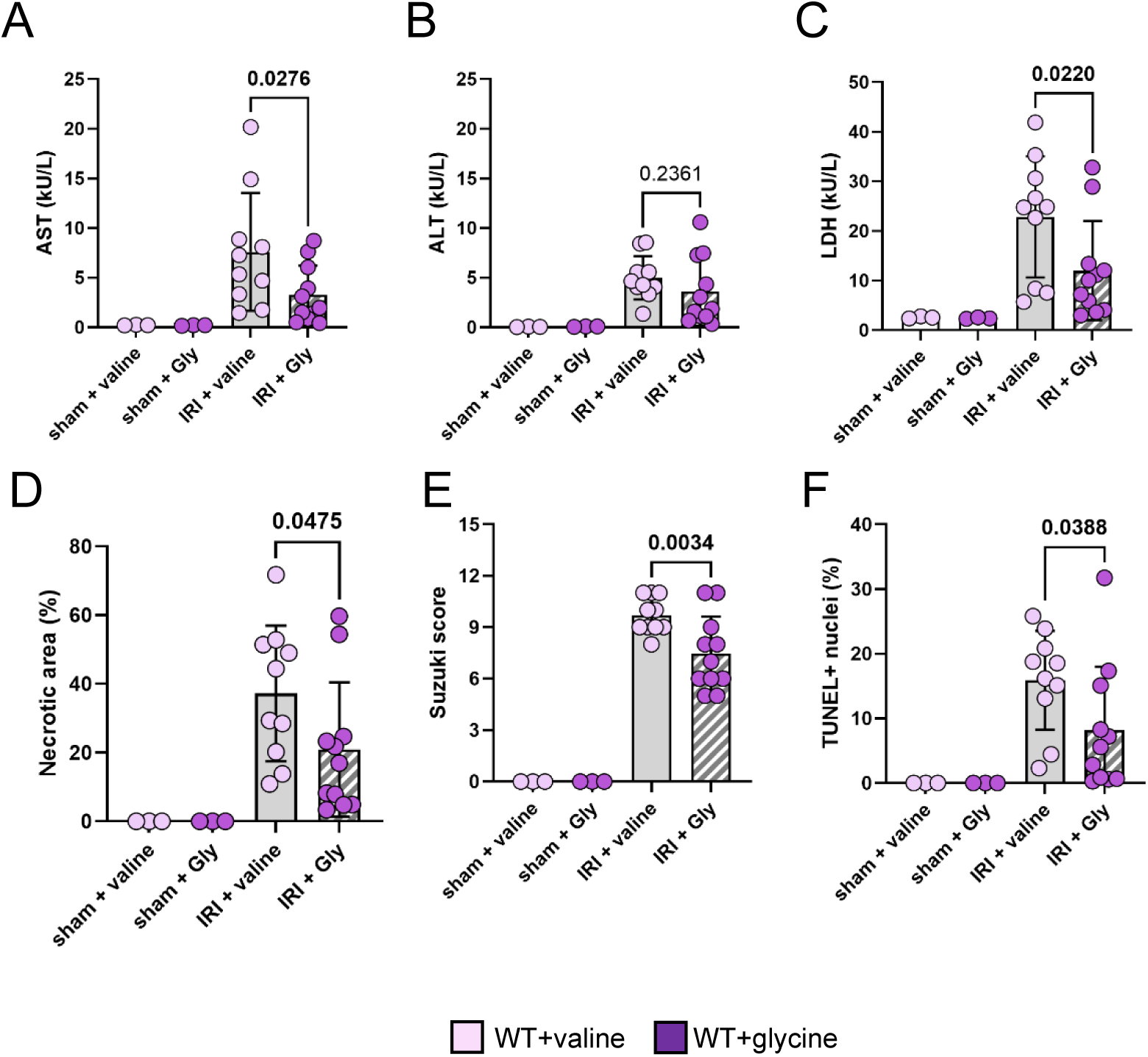
Glycine administration protects against liver ischemia-reperfusion injury. Wild-type C57BL/6 mice were administered with 0.5 mg/kg of sterile glycine or valine in PBS intraperitoneal 1 hour prior to ischemia and 0.5 mg/kg glycine or valine immediately following ischemia, followed by 6 hours of reperfusion. Group sizes: Sham valine, N = 3; Sham glycine, N = 3, IRI valine, N = 10; and IRI glycine, N = 11. (**A-C**) Hepatocyte-specific injury was assayed by serum AST and ALT, and general cellular cytotoxicity was measured by serum LDH. (**D-F**) Liver tissue analysis of the percent necrotic area, Suzuki score and the percentage of TUNEL-positive nuclei in the total area of the left lateral lobe and the left and right median lobes of the liver. Data points represent individual animals with the mean and standard error of the mean superimposed. *P*-values were determined by ANOVA with Šidák’s multiple comparison correction.

### Both macrophage and hepatocyte NINJ1 contribute to mouse IRI

Our data show that both Kupffer cells and hepatocytes express NINJ1 (**Figures 1** and **2**). Kupffer cells, the primary resident hepatic macrophages, make up 15-20% of normal murine livers (41) and are known to mediate hepatic IRI (42, 43). While hepatocytes make up 50-70% of the cells in the liver (41), the role of hepatocyte rupture in propagating IRI remains undefined. Our genetic knockout (**Figures 3** and **4**) and pharmacologic (**Figure 5**) approaches do not distinguish in which cell populations NINJ1 activation contributes to IRI. To delineate which NINJ1-expressing cell type(s) drive our observed protective phenotype in the whole animal KO models (**Figures 2** and **3**), we employed a conditional *Ninj1* knockout mouse (25). We targeted NINJ1 in the Kupffer cell and hepatocyte populations using *Ninj1* floxed mice crossed with either LysM-cre or Alb-cre, respectively, which were compared to littermate WT controls (*Ninj1* flox^+/+^ Cre^-/-^). Specific deletion of *Ninj1* in either macrophages or hepatocytes decreased serum markers (**Figures 6A-C** and **7A-C**) and histological evidence of IRI (**Figure 6D-F** and **7D-F**). Of note, hepatocyte-specific *Ninj1* deletion did not limit the extent of confluent necrosis or TUNEL positive cells following IRI (**Figure 7D,F**).

**Figure 6.**
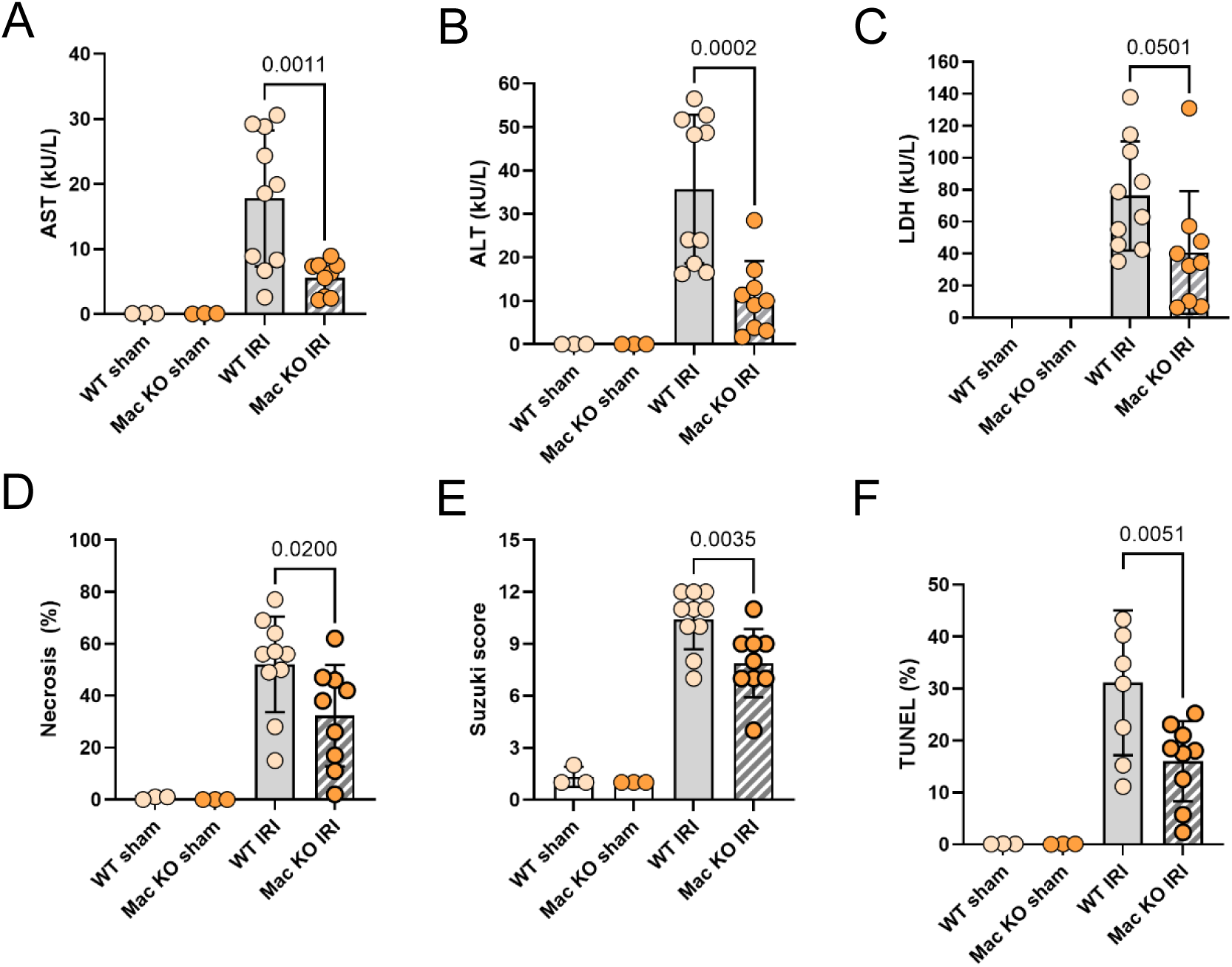
Macrophage-specific *Ninj1* knockout protects against liver ischemia-reperfusion injury in mice. Mixed-sex cohorts of macrophage-specific *Ninj1* knockout mice or their littermate controls underwent ischemia-reperfusion injury (IRI) consisting of 1 hour of 70% segmental warm ischemia followed by 6 hours of reperfusion prior to tissue collection. Sham laparotomy was used as a negative control. Group sizes: Sham WT, N = 3; Sham cKO, N = 3, IRI WT, N = 10; and IRI cKO, N = 9. (**A-C**) Hepatocyte-specific injury was assayed by serum AST and ALT, and general cellular cytotoxicity was measured by serum LDH. LDH data for sham-operated animals is unavailable due to inadequate sample for analysis. (**D-F**) Liver tissue analysis of the percent necrotic area, Suzuki score and the percentage of TUNEL-positive nuclei in the total area of the left lateral lobe and the left and right median lobes of the liver. Data points represent individual animals with the mean and standard error of the mean superimposed. *P*-values were determined by ANOVA with Šidák’s multiple comparison correction.

**Figure 7.**
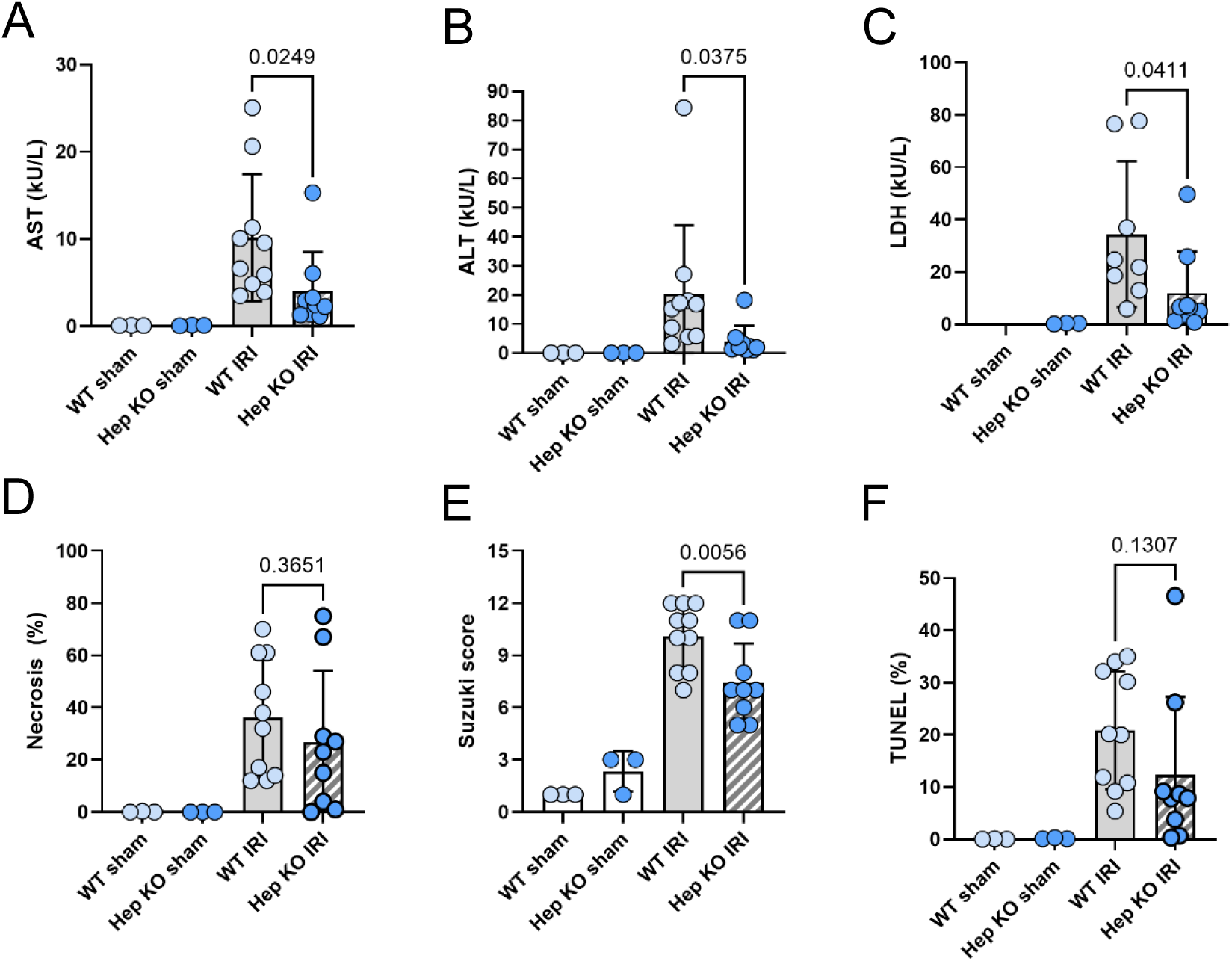
Hepatocyte-specific *Ninj1* knockout protects against liver ischemia-reperfusion injury in mice. Mixed-sex cohorts of hepatocyte-specific *Ninj1* knockout mice or their littermate controls underwent ischemia-reperfusion injury (IRI) consisting of 1 hour of 70% segmental warm ischemia followed by 6 hours of reperfusion prior to tissue collection. Sham laparotomy was used as a negative control. Group sizes: Sham WT, N = 3; Sham cKO, N = 3, IRI WT, N = 10; and IRI cKO, N = 9. (**A-C**) Hepatocyte-specific injury was assayed by serum AST and ALT, and general cellular cytotoxicity was measured by serum LDH. LDH data for WT sham-operated animals is unavailable due to inadequate sample for analysis. (**D-F**) Liver tissue analysis of the percent necrotic area, Suzuki score and the percentage of TUNEL-positive nuclei in the total area of the left lateral lobe and the left and right median lobes of the liver. Data points represent individual animals with the mean and standard error of the mean superimposed. *P*-values were determined by ANOVA with Šidák’s multiple comparison correction.

### Primary murine Kupffer cells and hepatocytes undergo NINJ1-mediated plasma membrane rupture in an *in vitro* hypoxia-reoxygenation model

Our *in vivo* data implicate both hepatocytes and Kupffer NINJ1 in hepatic IRI. To determine whether Kupffer cells are susceptible to NINJ1-mediated lytic cell death, primary Kupffer cells were isolated from the livers of *Ninj1^-/-^* and WT mice (**Figure 8A**). We confirmed that these cells were protected from plasma membrane rupture during pyroptosis when genetically devoid of *Ninj1*. Pyroptosis was induced by priming the cells with lipopolysaccharide (LPS) followed by treatment with the NLRP3 inflammasome-activating ionophore nigericin. Pyroptosis induction led to membrane rupture as quantified by LDH release (**Figure 8B**). LPS alone did not induce cell rupture, whereas LPS plus nigericin induced cell rupture in WT Kupffer cells, which was significantly decreased in *Ninj1^-/-^* Kupffer cells (**Figure 8B**). The magnitude of this effect was comparable to the pre-treatment of WT Kupffer cells with the cytoprotective agent glycine. Glycine did not confer additional protection to *Ninj1* KO Kupffer cells (**Figure 8B**), consistent with the concept that glycine is acting at the level of NINJ1 (26).

**Figure 8.**
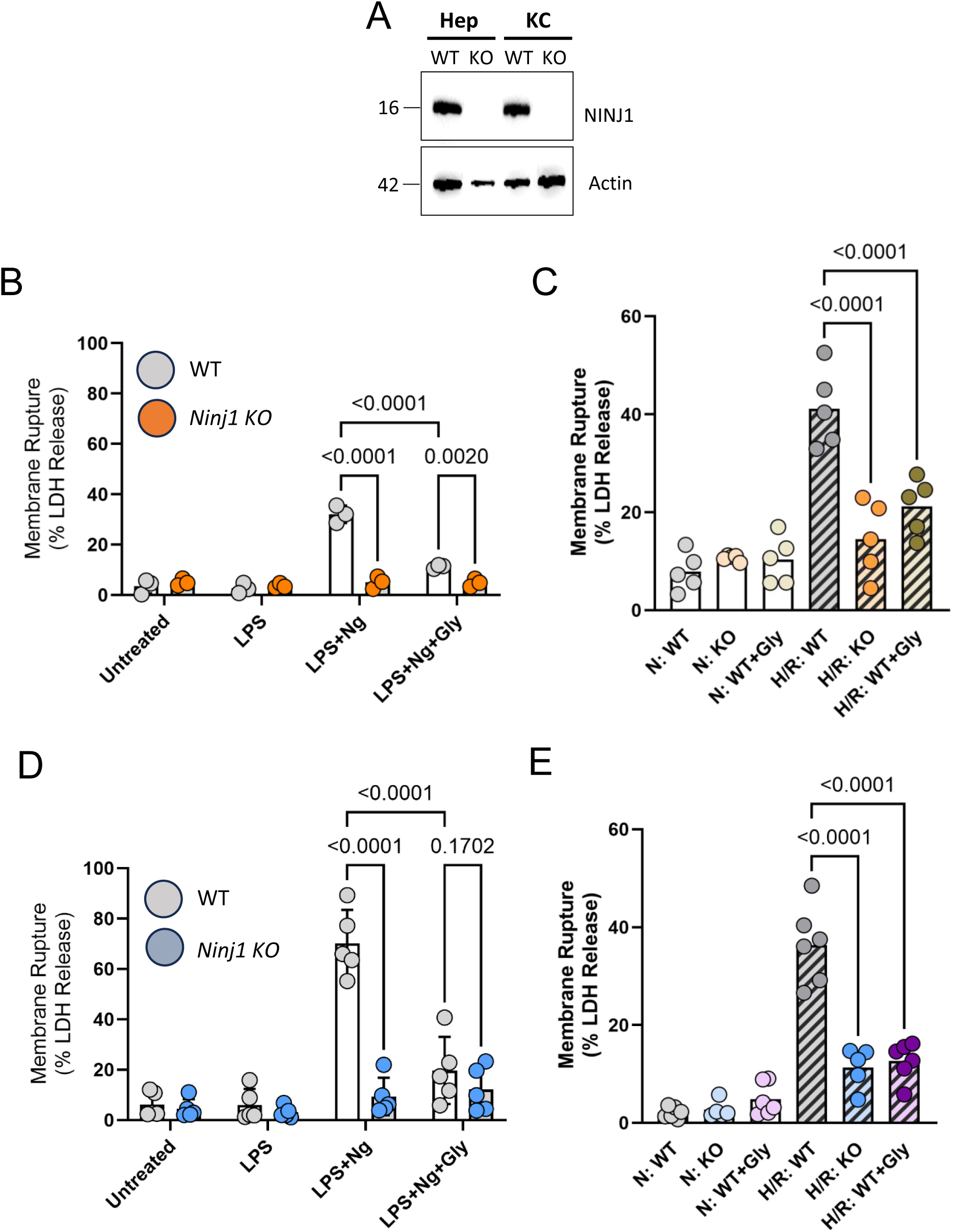
NINJ1 mediates plasma membrane rupture in hepatocytes and macrophages. (**A**) Primary hepatocytes and Kupffer cells were purified from wild-type and *Ninj1* knockout mice. NINJ1 expression was confirmed by Wester blot. β-Actin is shown as a protein loading control. (**B-E**) Pyroptosis was induced by priming with LPS (3 h) followed by treatment with nigericin (2 h). Untreated and LPS-alone are shown as controls. Hypoxia-reoxygenation injury was induced by incubating the cells in 1% O_2_ for 1 hour followed by normoxic conditions (21% O_2_) for 5 hours. Normoxia for 6 hours served as the control. Where indicated, cells were co-treated with glycine (5 mM). Membrane rupture was measured by LDH release. (**B-C**) Kupffer cells underwent (**B**) pyroptosis and (**C**) hypoxia–reoxygenation injury. In both models, glycine (5 mM) significantly reduced LDH release. *Ninj1* knockout Kupffer cells were protected from hypoxia-induced membrane rupture. (**D-E**) Hepatocytes underwent (**D**) pyroptosis and (**E**) hypoxia–reoxygenation injury. Glycine treatment (5 mM) decreased LDH release in wild-type cells in both contexts, while *Ninj1* knockout hepatocytes showed resistance to membrane rupture under hypoxia–reoxygenation. Data points represent independent experiments from cells derived from different animals with the mean and standard error of the mean superimposed. *P*-values were determined by ANOVA with Šidák’s multiple comparison correction.

To further delineate the mechanistic basis of NINJ1 activity in the Kupffer cell during IRI, we turned to an *in vitro* model of IRI consisting of hypoxic culture (1 hour at 1% O_2_) followed by normoxia for 5 hours (44). Hypoxia-reoxygenation induced lytic cell death in WT Kupffer cells, which was significantly reduced in *Ninj1^-/-^* cells (**Figure 8C**). Glycine treatment yielded a similar degree of cytoprotection in WT Kupffer cells as compared to the vehicle-treated condition and offered no additional protection to the *Ninj1* KO cells (**Figure 8C**).

To determine whether hepatocytes can similarly undergo active NINJ1-mediated plasma membrane rupture *in vitro*, primary hepatocytes were isolated from *Ninj1^-/-^* and WT littermates (**Figure 8A**). The cells were cultured in the presence of no stimulation, LPS alone, LPS plus nigericin or LPS plus nigericin and glycine. LPS alone did not induce lytic cell death, but LPS with nigericin induced pyroptosis in approximately 70% of cells (**Figure 8D**). Hepatocytes isolated from *Ninj1^-/-^* mice were resistant to plasma membrane rupture and glycine pre-treatment decreased LDH release from WT hepatocytes (**Figure 8D**). Glycine treatment of WT hepatocytes had a comparable effect to *Ninj1^-/-^* cells (**Figure 8D**). A similar phenomenon was noted utilizing hypoxia-reoxygenation injury, which induced a greater than 30 % release of cellular LDH from WT hepatocytes (**Figure 8E**). *Ninj1^-/-^* hepatocytes had substantial protection against this injury, and glycine was able to phenocopy this degree of protection in WT hepatocytes. To confirm that the cellular protection is due strictly to NINJ1 inhibition, the activation of caspase-1, a protease upstream of NINJ1 induced in pyroptosis, we utilized a fluorochrome labeled inhibitor of caspase-1 (FLICA) assay. Hypoxia-reoxygenation induced caspase-1 cleavage in hepatocytes), which was not prevented by glycine pre-treatment (**Supplemental Figure 5**). This demonstrates that the effect of NINJ1 inhibition does not disrupt upstream elements of the pyroptosis pathway.

Our *in vitro* data indicate that both Kupffer cells and hepatocytes express NINJ1, which executes membrane rupture in response to hypoxia-reoxygenation. These findings are consistent with the protection observed in our cell type-specific animals (**Figures 6** and **7**).

## Discussion

IRI of the liver allograft during transplant is associated with patient morbidity and mortality and worse overall outcome. Lytic cell death is a primary driver of the acute injury process, with NINJ1 positioned as the essential terminal effector of plasma membrane rupture (17). We previously demonstrated that NINJ1 inhibition with a neutralizing antibody abrogated several *in vivo* models of acute liver injury, including diminishing plasma markers rather than histopathologic indicators of IRI-induced hepatocellular injury in mice (25). In the current study, we address key knowledge gaps that remain. We first addressed the relevance of NINJ1 to human liver biology in the context of transplantation. By single cell RNA sequencing, spatial transcriptomics, and RNA-scope, *NINJ1* expression was predominantly seen within hepatocyte and Kupffer cell populations. Examining post-reperfusion samples of transplanted human livers used for transplant revealed increased NINJ1 activation in patients with evidence of severe early allograft dysfunction.

Utilizing *Ninj1^-/-^* knockout mice and rats, we established that NINJ1 deletion results in markedly reduced serum markers of hepatocellular injury and inflammation, tissue necrosis and cell death following IRI. Pharmacologic inhibition of NINJ1 with glycine phenocopies NINJ1 deletion *in vivo* and *in vitro*. In the murine liver, NINJ1 is similarly expressed by Kupffer cells and hepatocytes and mediates plasma membrane rupture in both cell types in response to cytotoxic signals, including hypoxia-reoxygenation. *In vivo*, both hepatocyte- and macrophage-specific deletion of *Ninj1* decrease IRI, suggesting both cell types play a role. Overall, these findings reveal that NINJ1 plays a crucial role in mediating plasma membrane rupture during IRI across cell types and species, offering a potential therapeutic target to promote organ health in liver transplant.

Our *in vivo* model of IRI employs a 1-hour period of ischemia followed by an acute period of 6-hours of reperfusion. With these conditions, the hepatocytes compartment experiences a significant injury as evinced by the significant increases in serum markers of hepatocyte injury. A recent study using bone marrow adoptive transfer to generate chimeric animals in which *Ninj1* is knocked out of the bone marrow compartment evaluated the effect of 1 hour of liver ischemia and either 6 or 24 hours or reperfusion (45). *Ninj1* knockout in the bone marrow compartment alone did not affect IRI injury in their model after 24 hours of reperfusion (e.g., AST, ALT, tissue necrosis and TUNEL-positive nuclei), whereas we observed markedly reduced indices of liver injury in our whole-body *Ninj1^-/-^* and glycine-treated animals at 6 hours. Hu *et* al depleted macrophages prior to BM transplant by treating the mice with liposomal clodronate. Although this approach does deplete macrophages, including Kupffer cells, clodronate liposomes may primarily impact the neutrophil compartment (46, 47). Furthermore, hepatocytes retained NINJ1 expression, which may partially explain some of the differences in findings. Nevertheless, combined, our study and that of Hu et al. (45) suggest NINJ1 activity is relevant in different liver cell populations at different phases of the IRI insult. Hence, it will be important to evaluate the long-term impact of NINJ1 knockout or inhibition following IRI. This could be accomplished by using an orthotopic liver transplantation model in the *Ninj1^-/-^*rats to understand if inhibition of NINJ1 and thereby limiting the early inflammatory response during IRI will impact the chronic sequelae of liver transplantation, such as fibrosis and graft rejection. This model would similarly provide opportunity to delineate the importance of NINJ1 in the donor graft and recipient respectively.

Mechanistically, our findings are consistent with a model in which NINJ1 inhibition prevents plasma membrane rupture in liver cells terminally injured by the ischemia-reperfusion insult. Plasma membrane rupture results in the release of intracellular contents, including DAMPs. Although smaller DAMPs (e.g. ATP) can efflux through small membrane pores, larger ones (e.g. high mobility group box 1, HMGB1) require membrane rupture to be released into the extracellular environment (48, 49). DAMPs, such as HMGB1, are known to contribute to hepatic IRI including in the context of liver transplantation (50–53). In addition to proteins, mitochondrial DNA (mtDNA) has also been implicated as an important DAMP in sterile inflammatory responses (54). mtDNA levels correlate with donor liver injury and primary graft dysfunction (55–57). Importantly, our data indicate that targeting NINJ1 only inhibits cell rupture: upstream pathways, including caspase-1 activation in pyroptosis, remain intact. These upstream pathways produce their own distinct inflammatory mediators (e.g. IL-1 cytokines) whose release do not require membrane rupture. The release of these DAMPs through cellular rupture involves NINJ1 oligomerization into multimeric complexes that act to disrupt the plasma membrane (22–24). The activation trigger for NINJ1 polymerization, particularly within the injured liver, is not fully elucidated. Recent studies suggest that plasma membrane strain or cell swelling as seen during the cell death process initiates NINJ1 oligomerization (19, 58). How molecular and biophysical changes in the plasma membrane are transmitted to NINJ1 require further investigation

While we have focused our investigations on the role of NINJ1 in executing plasma membrane rupture during liver injury, NINJ1 was first described as a cell-cell adhesion molecule as mediated by a putative cell adhesion domain within its N terminus (20, 59). As a cell adhesion molecule, it is plausible that NINJ1 enables recruitment of immune cells to the injured tissue. In our mouse model, however, we did not observe any differences in neutrophil and macrophage cell numbers in the *Ninj1* knockout animals as compared with their wild-type counterparts. Nevertheless, our data do not preclude the possibility that the protective effect of NINJ1 inhibition is mediated in part by suppressing cell-cell adhesion and therefore immune cell recruitment. The extent to which NINJ1 serves as a cell adhesion molecule merits re-evaluation given the recent structural study that positions the NINJ1 N terminus within the intracellular space (30).

Liver transplantation is the only curative treatment for end-stage acute and chronic liver disease but there remains a critical shortage of suitable organs for transplant (60). IRI during liver transplantation is a major clinical challenge that contributes to early allograft dysfunction with increased recipient morbidity or mortality (61). Expanding the donor pool to include more marginal organs (steatotic livers, livers procured from older donors or from donation after cardiocirculatory death), increases the risk of severe IRI and early graft dysfunction, as these organs are at higher risk (61–63). Developing novel approaches to reduce IRI may help expand the donor pool. Our ability to link glycine’s cytoprotective effect to NINJ1 activation and protection against IRI has potential translational implications to this end. Glycine has long been known to suppress reperfusion injury when added to liver perfusates (39).

Alternatively, inhibition of NIN1 can be achieved via a blocking antibody (clone D1), which ameliorates acute liver injury across multiple mouse models, including ischemia reperfusion injury (25). Clone D1 pre-treatment significantly reduces circulating levels of hepatic transaminases six hours post-reperfusion injury although a significant difference in tissue necrosis was not observed (25). This difference in efficacy between the D1 pharmacologic intervention and our knockout studies may reflect incomplete inhibition by the D1 antibody due to the dosing regimen employed in our early study. Regardless, anti-NINJ1 therapies could be applied to liver perfusates during the ischemic phase of transplantation directly to the graft or via expanding utilization of normothermic machine perfusion technologies to enhance cellular integrity and thereby limit the extent of graft injury and dysfunction following transplantation.

## Methods

Our study examined male and female human specimens as well as male and female animals. Similar findings are reported for both sexes.

### Human liver samples

All human liver samples were collected with institutional ethics approval from the Research Ethics Board at the University Health Network (UHN), Toronto, Canada (REB# 22-5892) and The Hospital for Sick Children (REB# 1000082047). Liver tissue was obtained from the UHN Multi-Organ Transplant (MOT) Biobank (REB# CAPCR 15-9179) Biobank. All adult patients (age over 18 yr) who underwent liver transplantation at UHN from 2016-2022 utilizing allografts from a deceased donor for which liver biopsy cryopreserved tissue specimens are stored in the MOT biobank. Inclusion criteria for the present study were the availability of post-reperfusion biopsy liver tissue and either AST <500 or >5000 within the first 7 days post-transplant for the control and early allograft dysfunction cohorts, respectively (28, 64).

Patients that received a live donor liver allograft or for which available clinical data were incomplete were excluded. Convenience sampling of the first available 18 patients meeting inclusion criteria was used. Patient demographic and clinical characteristics are shown in **Supplemental Table 1**.

### Single cell RNA sequencing, spatial transcriptomics and RNA in situ hybridization

Single-cell RNA-sequencing (scRNA-seq) and Visium spatial transcriptomic data from non-diseased human donor livers were obtained from (65). Library preparation, sequencing, and primary bioinformatic processing for this dataset, including demultiplexing, alignment, UMI counting, quality control, clustering, and initial dimensionality reduction were performed as described in the original publication and were not repeated here. For the present study, gene–cell expression matrices and associated metadata from AnnData (.h5ad, HDF5-based) files were imported into R (version 4.5.1) using Seurat (version 5.3.1). The published cell annotations and precomputed low-dimensional embeddings were retained, and analysis was restricted to non-diseased donor samples. Uniform manifold approximation and projection (UMAP) coordinates from the original analysis were used directly for visualization of *NINJ1* expression. RNA-scope (Bio-Techne) was completed on paraffin-embedded human liver samples as per the Manufacturer’s instructions. In situ hybridization was completed using the following probes *CD68*-C1 probe #402681, *NINJ1-*C4 probe #540051-C4, *ALB*-C3 probe #600941-C3, and *VWF*-C2 probe #413401-C2. Images were acquired using Imaris acquisition software (Oxford Instruments; RRID: SCR_00737).

### Animals

All animal procedures were conducted under protocols approved by the Animal Care Committee at The Hospital for Sick Children and in accordance with animal care regulation and policies of the Canadian Council on Animal Care. Mice and rats were housed in same-sex cohorts in polycarbonate cages with *ad libitum* access to food and water. Housing rooms were temperature and humidity controlled with 14:10 hour light:dark cycles.

*Mice -- Ninj1* knockout (*Ninj1^-/-^*) mice on a C57BL/6 background were previously described (17) with *Ninj1* wild-type littermate mice used as controls. In glycine treatment studies, wild-type C57BL/6 animals were purchased from Jackson Laboratory (strain Cat. No.000664, Bar Harbour, MA, USA). Ninj1^fl/fl^ mice with a floxed exon 3 were described previously (25). Hepatocyte and macrophage-specific *Ninj1* knockout mice were generated by breeding *Ninj1^fl/fl^* mice with Albumin-cre (B6.Cg-Speer6-ps1^Tg(Alb-cre)21Mgn^/J; Jackson Laboratory, strain Cat. No.003574) and LyzM-cre (B6.129P2-Lyz2^tm1(cre)Ifo^/J, Jackson Laboratory, strain Cat. No.004781). Genotyping by PCR was conducted as per instructions provided by Jackson Laboratory.

*Rats --* The *Ninj1*− allele was generated by ENVIGO using established CRISPR methodology and electroporation of Cas9 in complex with sgRNAs into Sprague Dawley rat zygotes (Taconic). Two sgRNAs were designed to generate a 368bp deletion at genomic coordinates 12301-12668. NHEJ activity of the upstream sgRNA was detected using PCR primer set A (5’-tcttgctggagaccagtgtg and 5’-gaagaaggcgaactcattgc) yielding an expected band size of 396bp. NHEJ activity of the downstream sgRNA was detected using PCR primer set B (5’-gcaatgagttcgccttcttc and 5’-gagccacagggatctaagga) yielding an expected band size of 338bp. *Ninj1^-/-^* was genotyped with PCR primer set C (5’-tcttgctggagaccagtgtg and 5’-gagccacagggatctaagga) yielding ∼362bp *Ninj1^-/-^*. Wild-type rats were littermates. *Ninj1*

### Segmental ischemia-reperfusion injury model

Mixed-sex cohorts of 6- to 12-week-old mice or rats of either *Ninj1^-/-^*or wild-type littermate controls were used in a 70% segmental ischemia-reperfusion model (66). Under isoflurane anesthesia, a sagittal midline laparotomy was made, and an atraumatic vascular clamp placed on the portal vein and the hepatic artery to block blood flow to the left and medial lobes of the liver. After 1 hour, the clamps were removed and the animal was returned to its home cage to allow for reperfusion. Following 6 hrs of reperfusion, the animal was euthanized by cardiac puncture under general anesthesia and blood and tissues were collected for analysis. Sham laparotomy, where the vascular pedicle was exposed but not clamped, was used as a negative control. Where indicated, animals were randomly allocated into either the glycine-treatment group or the valine-treatment group. Animals were treated with intraperitoneal injection 1 hour before and immediately after removal of the vascular clamps with glycine (0.5 mL of 100 mM in PBS; Cat. No.G7126, MilliporeSigma, Burlington, MA, USA) or valine (0.5 mL of 100 mM in PBS, Cat. No. 94640, MilliporeSigma). If ischemia-reperfusion did not induce injury in both the left and medial lobes of the liver, these animals were excluded with the assumption the vascular clamping was not optimal. Each animal was defined as an experimental unit for the purpose of sample size determination. Our primary outcome was confluent necrosis as assayed by histopathology. Secondary outcomes included serum biochemistry for liver injury and cell death as assayed by TUNEL assay on pathology samples. Experimenters were not blinded to surgical intervention (sham *vs* IRI) but were blinded to genotype or pharmacologic treatment group allocation during the experiment and subsequent analyses. Our study adhered to the ARRIVE 2.0 guidelines for animal experimentation (67).

### Serum biochemistry

Blood was collected by cardiac puncture at the end of the reperfusion period and placed on ice. Serum was prepared by centrifugation of the clotted blood. A quantitative LDH colorimetric assay kit was used to measure LDH in the serum samples as per the manufacturer’s instructions (Cat. No. MAK006, MilliporeSigma). Serum AST and ALT levels were measured at the Pathology Core laboratory at The Centre for Phenogenomics (Toronto, Canada) using a Beckman Coulter AU480 clinical chemistry analyzer by photometry testing (Beckman Coulter Life Sciences, Indianapolis, IN, USA) in combination with appropriate calibrators (Beckman Coulter Lyophilized Chemistry Calibrator Levels 1 and 2) and quality control materials (Liquid Assayed Multiqual 1 and 3, Bio-Rad, Hercules CA, USA).

### Histology

All ischemic and reperfused liver lobes were collected, paraffin-embedded, and sectioned at a thickness of 4 µm. Serial sections were then stained with haematoxylin and eosin or terminal deoxynucleotidyltransferase dUTP nick-ending labeling (TUNEL; Terminal Transferase, Cat. No. 03 333 566 001 and Biotin-16-dUTP, Cat. No. 11 093 070 910, Roche Diagnostics, Inc., Indianapolis, IN, USA). Tissue specimen processing and staining were conducted at the Spatio-Temporal Targeting and Amplification of Radiation Response (STTARR) Innovation Centre (Toronto, Canada). Slides were imaged using a Panoramic Flash II Slide Scanner (3DHistech Inc., Budapest, Hungary) and visualized using HALO Image Analysis Platform (RRID: SCR_018350; Indica labs, Albuquerque, NM, USA). To evaluate the extent of necrosis within the collected liver samples, the DenseNet V2 classifier supervised machine learning algorithm (HALO Image Analysis Platform) was trained to recognize necrotic tissue in the H&E stains and applied to the entire liver sample. To evaluate the TUNEL-positive nuclei, we used the nuclei-seg pretrained classifier plug-in into the Multiplex IHC module to distinguish TUNEL-stained (brown) from Hematoxylin-stained (blue) nuclei. The software was trained on multiple samples prior to developing an effective generalized classifier. Negative annotation tool was used to exclude regions with high background staining on the edges or regions of poor tissue sample integrity (due to edge-effects or tears in the tissue).

### Immunohistochemistry of macrophage and neutrophils

Formalin-fixed paraffin-embedded tissue samples mounted on slides, stained with hematoxylin, were dewaxed and rehydrated using a standard protocol. Samples were also blocked for endogenous peroxidase activity in 3% H_2_O_2_. Antigen retrieval was performed in a citrate buffer (ab93678, Antigen Retrieval Buffer 100x pH 6.0, Abcam) with heating at 98°C. Following washes, samples were protein-blocked with Dako’s Serum Free Protein Block (X0909, Dako) followed by incubation with primary antibodies: rabbit anti-Ly6G (1:200 dilution, 87048, Cell signaling) or rabbit anti-F4/80 (1:100, Cat. No. 70076S, Cell Signaling) 1:100 in dako diluent (Cat. No.S0809, Dako), incubate for 1 hour at RT. After washes, samples were incubated with the biotinylated Anti-rabbit Ig, secondary antibody (BA-1000, Vector Labs) for 30 minutes at RT. Detect with Avidin biotin complex (ABC) system (Vector Labs VECTASTAIN® Elite® ABC-HRP Kit, Peroxidase (Standard) PK-6100) incubate for 30 minutes at RT Incubate in DAB (3,3′-diaminobenzidine; ab64238 DAB Substrate Kit, Abcam) solution made according to product datasheet for 10 minutes. After, sample dehydration through a graded alcohol series, the samples were cleared in xylene and mounted with coverslips. To quantify the number of neutrophils and macrophages within the mouse liver tissue samples, we used the Multiplex IHC module in HALO Image Analysis Platform to distinguish Ly6G-stained (neutrophils) and F4/80-positive (macrophages) cells.

### Cells isolation and culturing

*Primary macrophages-* Primary bone marrow derived macrophages (BMDM) were harvested, as previously described (26), from the femurs and tibia of mixed-sex cohorts of either wildtype or *Ninj1* knockout mice on a C57BL/6 background. In brief, the bones were cleaned, the ends cut, and centrifuged to collect bone marrow into sterile PBS. Following a wash in PBS, the cells were plated in DMEM with 10 ng/mL M-CSF (Cat. No. 315-02; Peprotech Inc, Cranbury, NJ, USA). After 5 days of culture, the BMDM were detached from the dishes with TBS with 5 mM EDTA, resuspended in fresh DMEM and plated.

*Primary hepatocytes and Kupffer cells-* Primary mouse hepatocytes and Kupffer cells were isolated as described (68). In brief, under general anesthesia (2 g/kg urethane), the inferior vena cava was cannulated, and the liver perfused to remove blood and chelate calcium using EDTA (30 ml EDTA 0.5 mM in PBS at 300 ml/h) before administering a Liberase collagenase digestion solution (20 ml liberase 45 μg/ml (LIBTM-RO, Liberase™ TM Research Grade Roche Diagnostics, Indianopolis, IN, USA) in HBSS (Cat. No. 311-515-CL, Wisent Bioproducts, St-Bruno, QC, Canada) at 300 ml/h) maintained at 37°C, to dissociate the extracellular matrix. In a bio-safety cabinet, the liver was dissected in a 10 cm dish and cells were rendered in suspension using tweezers in HBSS then filtered through a 40 μm cell strainer and collected at 50 x g, 2 min. Mainly hepatocytes resided in the pellet and Kupffer cells remain in the supernatant and were purified using density-based separation on a 36% (v/v) Percoll gradient (Cat. No. 17089101, Cytiva, Marlborough MA, USA). Hepatocytes were seeded on 1% gelatin-coated plates in DMEM-high glucose supplemented with 2 mM L-glutamine, pyruvate (Cat. No. 319-0007-CL, Wisent Bioproducts) with addition of 10% FBS and 1% Pen-Strep and for 3 hours, then the media was changed for overnight in William E medium supplemented (Cat. No. W4125, MilliporeSigma, Burlington, ME, USA) with 10% FBS, 1.5 g/L sodium bicarbonate, 1% Pen-Strep. Kupffer cells were plated on gelatin-precoated plates with the same DMEM-high glucose media used for hepatocytes, that was also supplemented with 10 ng/ml of M-CSF. Primary cells were used within 48 hours of isolation.

### Pyroptosis induction

Hepatocytes were grown in DMEM low glucose with 0.5% FBS and primed with (0.5 µg/ml) LPS from E. coli serotype 055:B5 (Cat. No. L5418, MilliporeSigma), which was reconstituted at a stock concentration of 1 mg/ml. After 3 h of LPS priming, pyroptosis was induced by addition of 20 µM nigericin (Cat. No. N7143; stock 10 mM in ethanol, MilliporeSigma) for 120 min, as indicated. Where indicated, cells were treated with 5 mM glycine (Cat. No. G7126, MilliporeSigma) or 5 mM valine (Cat. No. 94640, MilliporeSigma) following the 3 h of LPS priming and prior to treatment with nigericin. Kupffer cells were cultured under identical media conditions and primed with 0.5 µg/ml LPS for 3 h. Pyroptosis was subsequently induced by treatment with 20 µM nigericin for 30 min. As described for hepatocytes, where indicated, Kupffer cells were incubated with 5 mM glycine or 5 mM valine immediately after the 3 h LPS priming period and included with the nigericin treatment.

### Hypoxia-reoxygenation treatment

Hypoxia-reoxygenation was performed in primary cells (hepatocytes or Kuffer cells) grown in 12-well plates with DMEM low glucose with 0.1% FBS and pen/strep antibiotics based on the published protocol with modifications (44). Cells received a media replacement with 1% O2 pre-incubated media, for a 1-hour incubation in a Oxycycler^TM^ (Biospherix Ltd., Parish NY, USA) with a 1% O2 / CO2 atmosphere.

Afterwards, the cells were switched to 21% O2 pre-incubated media (DMEM + 0.1% FBS, pen/strep) and incubated for 5 hours in a 21% O2 / CO2 normal atmosphere incubator with or without glycine (5 mM) supplementation.

### In vitro LDH release assay

Cells were seeded at 200,000 cells per well in 12-well plates. Following treatment, cell culture supernatants were collected and centrifuged for 5 minutes at 500 x g to eliminate cellular debris. On ice, the cell plates were washed once with cold PBS and lysed in lysis buffer provided in the LDH assay kit (Cat. No. C20300, Thermo Fisher Scientific, Waltham, MA, USA) containing protease inhibitors (Pierce tablet, Cat. No. A32955, Thermo Fisher Scientific). The collected supernatants and lysates were assayed for LDH using a colorimetric assay kit following the manufacturer’s instructions.

### Fluorescence microscopy

*NINJ1 immunofluorescence*- Primary hepatocytes from *Ninj1* wildtype and *Ninj1* knockout mice were harvested as described above and cultured on glass coverslips. Cells were washed with PBS and fixed in 4% paraformaldehyde (Cat. No. 15710, 16% stock, Electron Microscope Sciences, Hatfield, PA, USA) in PBS at room temperature for 15 min. The cells were washed 3 times with PBS then blocked in 10% donkey serum in PBS for 1 hour. Cells were then incubated overnight at 4 °C in rabbit monoclonal anti-mouse NINJ1 primary antibody (Clone 25, kindly provided by Dr. Vishva Dixit, Genentech Inc. South San Francisco, CA, USA) at 10 µg/mL. The cells were washed three times with PBS and incubated in Cy3-conjugated donkey anti-rabbit secondary antibody (1:1000 dilution, Cat. No. 711-165-152, Jackson ImmunoResearch) in PBS supplemented with 1% donkey serum for 1 hour at RT. Following incubation, secondary antibodies were removed and the nuclei labeled with DAPI at 0.5µg/mL for 5 min. Cells were then washed with PBS before being imaged by spinning disk confocal microscopy (Quorum Technologies Inc. Puslinch, Canada) on a Zeiss Axiovert 200M microscope with a 63× objective. Images were acquired by a CCD camera (Hamamatsu Photonics, Bridgewater, NJ, USA) driven by the Volocity software (RRID: SCR_002668, Quorum Technologies Inc. Puslinch, Canada). Three images per treatment per experiment were taken and images were processed using ImageJ.

*FAM-YVAD-FMK fluorescence*- Caspase 1 activation was measured by confocal microscopy using the Pyroptosis FAM Caspase-1 Kit (Cat. No. ICT9145, Bio-Rad Hercules, CA, USA) The cell-permeant fluorescent probe, FAM-YVAD-FMK, labels active capase-1 in live cells through the specific binding of the YVAD peptide to caspase-1 and irreversible covalent bond formation by the fluoromethyl ketone (FMK) reactive group. Excess unreacted reagent is washed out and fluorescent signal of the Fluorescein (FAM) remaining in the cells is proportional to the caspase 1 activation. Hepatocytes were seeded on 18mm glass coverslips in 12-well plates at 1 x 10^5^ cells per well and treated with hypoxia-reoxygenation as described above. FAM-YVAD-FMK was added to the cells for 60 min at the working dilution, and then the cells were incubated with fixative provided with the kit, following the manufacturer’s instructions. After staining, coverslips were mounted onto glass slides using ProLong diamond antifade mounting medium (Cat. No. P36965, Thermo Fisher Scientific). Cell images were collected, processed and quantified as described above.

### SDS-PAGE and Western blotting

Cell grown in 6-well plates, treated as indicated, were washed rapidly with PBS and lysed with 200 μl RIPA containing protease inhibitors (50 mM Tris HCl, 150 mM NaCl, 1.0% (v/v) NP-40, 0.25% (w/v) Sodium Deoxycholate, 1.0 mM EDTA, 0.1% (w/v) SDS and 0.01% (w/v) sodium azide at a pH of 7.4 supplemented with Pierce™, protease inhibitor tablet (Cat. No. A32955, Thermo Fisher Scientific).

Lysates were centrifugated at 14,000 × g for 30 min and the supernatants were saved. The protein concentrations of cell lysates were measured using the Pierce™ BCA Protein Assay Kit (Cat. No. 23227, Thermo Fisher Scientific). The lysates were adjusted to equal protein concentrations and were mixed 1:1 with 2× Laemmli sample buffer and incubated for 5 min at 95 °C. Samples (10 μg protein) were resolved using SDS-PAGE, 12 or 15% polyacrylamide gels, transferred to Immun-Blot PVDF Membrane (Cat. No. 1620177, Bio-Rad), blocked with 1% BSA in TBS-Tween (150 mM NaCl, 0.1% (w/v) Tween® 20 detergent, 20 mM Tris-HCl, pH 7.5) then incubated with primary antibodies indicated according to the manufacturer’s or published recommendations. Antibodies used: anti-caspase-1 (p20) (1:1000, Cat. No. AG-20B-0042-C100, AdipoGen Life Sciences, Inc. San Diego, CA, USA); Anti-NINJ1 rabbit monoclonal (clone 25, 1 μg/ml) antibody (17); Anti-β-Actin mouse monoclonal (1:1000, Cat. No. A1978, MilliporeSigma). Following incubation with the primary antibodies, PVDF membranes were washed with TBS-Tween, and incubated with Goat-anti-rabbit or Donkey-anti-mouse horseradish peroxidase-linked secondary antibody (Jackson ImmunoResearch), and washed again before visualizing via reaction with the Novex™ ECL Chemiluminescent Substrate Reagent Kit (Cat. No. WP20005, Thermo Fisher Scientific) and using ChemiDoc XRS+ Imaging System (Bio-Rad).

### Blue-Native PAGE and Western blotting

*Cells*- Bone marrow-derived macrophages were seeded in 12-well plates. The following day the cells were treated as indicated in the figure legends, then placed on ice. The cells were washed with PBS then lysed in ice-cold digitonin lysis buffer (150 mM NaCl, 150 mM Tris-HCl, pH 7.5, 1% digitonin supplemented with Pierce™, protease inhibitor tablet) scraped into microtubes, and incubated 10 min on ice with periodic mixing. Lysates were centrifuged at 20,800 x g for 30 min and the supernatant was collected and protein concentration determined using the DC protein assay kit (Bio-Rad, Cat. No. 5000111). Equal amounts of protein/ condition were mixed with Native-PAGE sample buffer prior to being resolved using NativePAGE 3-12% Bis-Tris gels (Cat. No. BN1001BOX, Thermo Fisher Scientific) according to the Manufacturer’s instructions.

*Tissue-* Liver tissue lysates were prepared as previously described by (69) with some modifications. Briefly, 20 mg of liver tissue were homogenized in 500 μl of sucrose buffer (250 mM sucrose, 20 mM imidazole/HCl pH 7.0, supplemented with Pierce™, protease inhibitor tablet) using a Teflon Potter-Elvehjem homogenizer for 20 strokes, then centrifuged for 720 x g for 5 min to remove unhomogenized material and nuclei. Supernatants were then centrifuged at 20,800 x g for 30 min to collect a crude plasma membrane fraction. The pellet was resuspended in 60 μl of solubilization buffer (50 mM NaCl, 50 mM Imidazole, 2 mM 6-aminohexanoic acid, 1 mM EDTA, pH 7.0) and 25 μl was retained for SDS-PAGE (see below) and to the remaining 35 μl, 20 μl of 20% digitonin was added and the samples were incubated 10 min on ice with periodic mixing. Samples were then centrifuged for 30 min at 20,800 x g and supernatants were retained, and a small 2μl aliquot was removed (and diluted) for protein determination using the *DC* Protein Assay Kit (Cat. No. 5000111, Bio-Rad). Next, 5 μl of 50% glycerol and 10 μl of 5% w/v Coomassie G-250 were added to the samples. 10 μg protein samples were resolved on NativePAGE 3-12% Bis-Tris gels using an XCell SureLock™ Mini-Cell electrophoresis system (EI0001,

Thermo Fisher Scientific) according to the Manufacturer’s instructions. In one of the gel lanes, the NativeMark™ Unstained Protein Standard (LC0725, Thermo Fisher Scientific) was loaded. After performing the electrophoresis, the gels were electro-transferred to PVDF membranes and were Western blotted (as described above) with anti-NINJ1 rabbit monoclonal (clone 25) antibody at 10 μg/ml. For SDS-PAGE experiments to assess total NINJ1 in the homogenized liver tissue from above, an aliquot was used to determine protein concentration (with the Pierce™ BCA Protein Assay Kit (Cat. No. 23227, Thermo Fisher Scientific)) and 10 ug samples were mixed 1:1 with 2× Laemmli sample buffer and separated on a 12% SDS-PAGE gel and Western blotted with anti-NINJ1 rabbit monoclonal antibody (clone 25, 1 μg/ml) and anti-β-Actin (1:1000) as described in the Western blotting protocol.

### Quantification and statistics

Statistical testing was calculated using Prism 9.0 (GraphPad Software Inc, La Jolla, CA; RRID:SCR_002798). Data are provided as mean ± SEM. Groups were compared using Student t test for two groups and ANOVA for three or more groups with Šidák’s test for multiple comparison. All collected data was analyzed and a *P* value < 0.05 was considered statistically significant. For non-quantitative data (e.g., western blots), experiments were replicated three times with representative images provided in the figures.

## Data availability

All data are depicted in the main text or the supplementary materials. Requests for raw data and materials are available upon request to B.A.S.

## Acknowledgements

We thank Dr Vishva Dixit (Genentech, Inc) for providing the anti- NINJ1 antibodies and the *Ninj1* knockout mice. We thank Dr Jason Maynes (Hospital for Sick Children) for use of his Oxycycler^TM^ (Biospherix Ltd). Graphical schematics and Figure layout were produced with BioRender (RRID:SCR_018361) and Inkscape (RRID:SCR_014479), respectively.

## Author contributions

Conceptualization: JM, BM, NMG, BES, BAS

Methodology: JM, BM, PJB, IBS, NK, SM, SAF, NMG, BES, BAS

Investigation: JM, BM, YZ, FA, DT, CJH, AV, GY, DMA, AM, IS, GG, PJB, IBS

Visualization: JM, BM, FA, DT, CJH, BES, BAS

Funding acquisition: NMG, BES, BAS

Project administration: BES, BAS

Supervision: NK, SAF, NMG, BES, BAS

Writing – original draft: YZ, PJB, BES, BAS

Writing – review & editing: JM, BM, YZ, FA, DT, CJH, AV, GY, PJB, IBS, NK, NMG

JM and BM are co-first authors. Their listed order is reverse alphabetical.

BES and BAS are co-senior authors. Their listed order is reverse alphabetical.

---

**Supplemental Figure 1.**
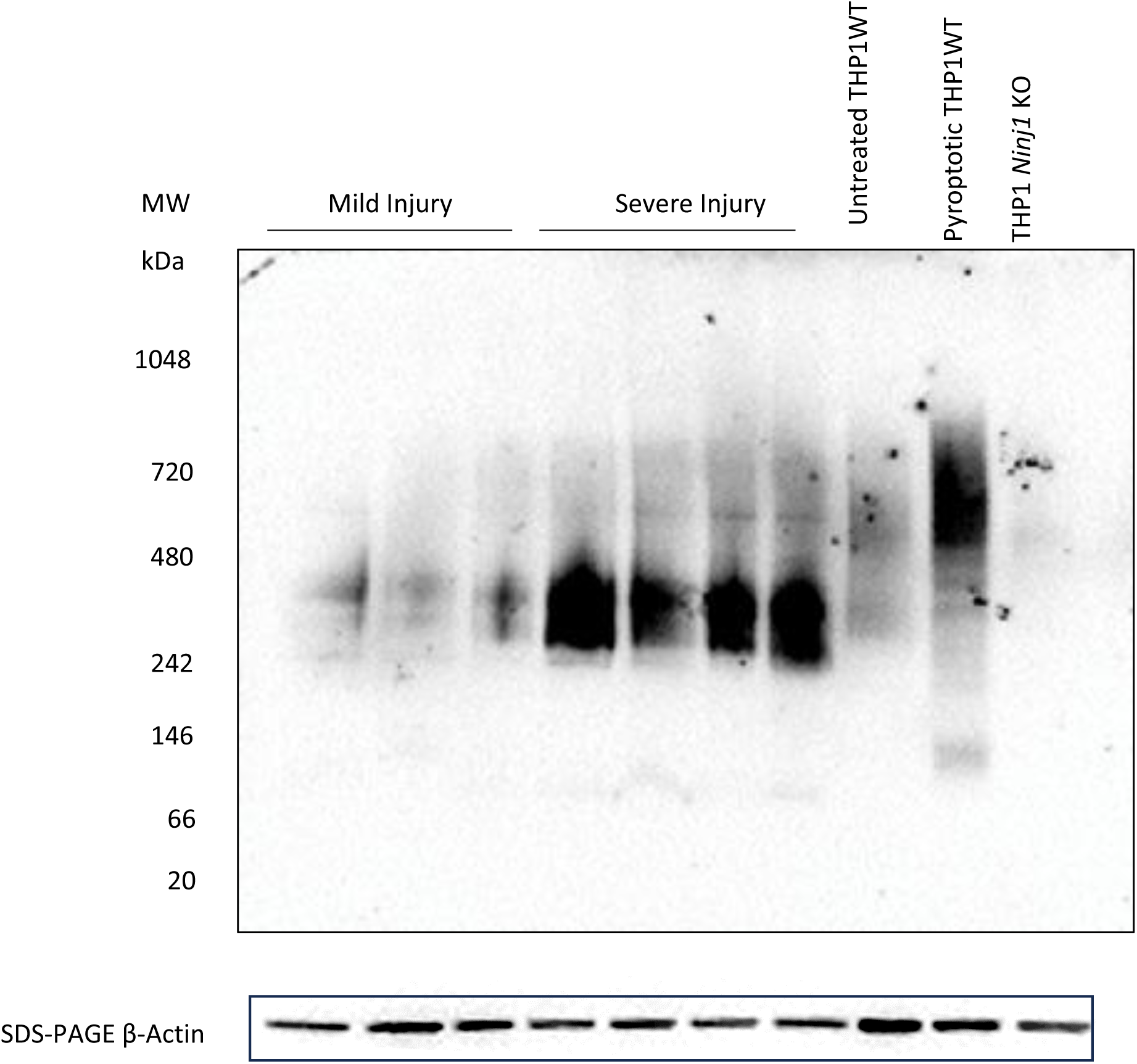
Representative blue native-PAGE of human NINJ1 in patient liver samples from low and high early allograft dysfunction groups. Human liver samples from patients with mild injury (lanes 1–3) and severe injury (lanes 4–7; defined by AST > 5000 U/L) were subjected to BN-PAGE to assess NINJ1 clustering. Increased high–molecular weight NINJ1 complexes (≈242–480 kDa) were predominantly observed in the severe injury group. THP1 wild-type cell lysates (untreated), THP1 wild-type cell lysates under pyroptosis (priming with 0.5 µg/ml LPS from E. coli serotype 055:B5 for 3 hours followed by 20 µM nigericin treatment for 2 hours), and THP1 *NINJ1* knockout cell lysates were included as controls. Corresponding SDS-PAGE immunoblots for β-actin confirm similar protein loading.

**Supplemental Figure 2.**
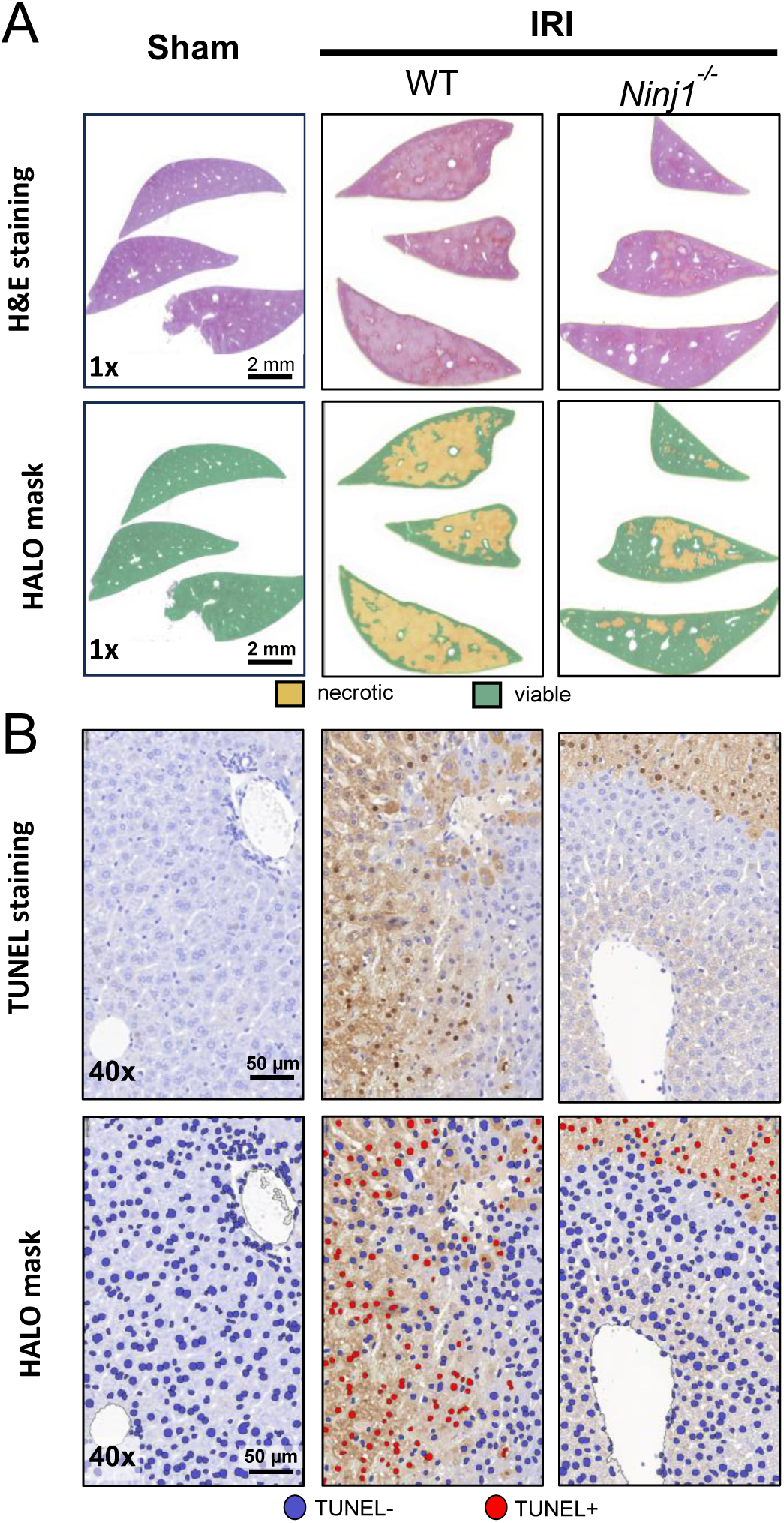
Examples of automated classification of necrotic liver tissue and TUNEL positive cells. As described for Figure 3, mixed-sex cohorts of *Ninj1* knockout mice or their littermate controls underwent ischemia-reperfusion injury (IRI) consisting of 1 hour of 70% segmental warm ischemia followed by 6 hours of reperfusion prior to tissue collection. Sham laparotomy was used as a negative control. Group sizes: Sham WT, N = 6; Sham KO, N = 4, IRI WT, N = 15; and IRI KO, N = 8. (**A**) To evaluate the extent of necrosis within the collected liver samples, the DenseNet V2 classifier supervised machine learning algorithm (HALO Image Analysis Platform) was trained to recognize necrotic tissue in the H&E stains and applied to the entire liver sample. Representative images of the H&E (*top*) and classified histology (*bottom*) are shown. (**B**) To evaluate the TUNEL-positive nuclei, we used the nuclei-seg pretrained classifier plug-in into the Multiplex IHC module to distinguish TUNEL-stained (brown) from Hematoxylin-stained (blue) nuclei. The software was trained on multiple samples prior to developing an effective generalized classifier. Representative images of the TUNEL stain (*top*) and classified nuclei (*bottom*) are shown. TUNEL negative and positive nuclei are shown in blue and red, respectively. Scale bars are as shown.

**Supplemental Figure 3.**
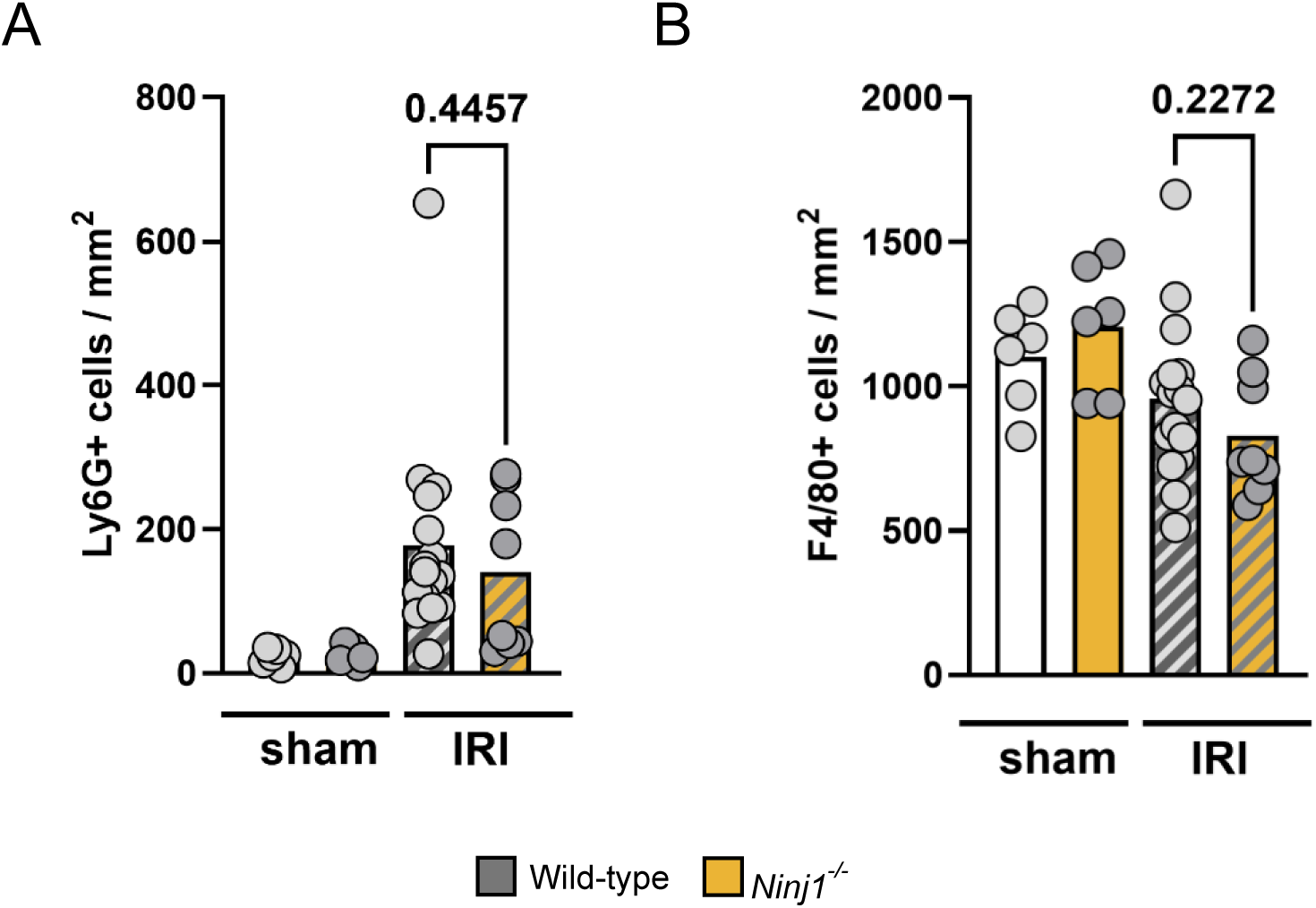
Macrophage and neutrophil recruitment to the liver following ischemia-reperfusion injury in mice. As described for Figure 3, mixed-sex cohorts of *Ninj1* knockout mice or their littermate controls underwent ischemia-reperfusion injury (IRI) consisting of 1 hour of 70% segmental warm ischemia followed by 6 hours of reperfusion prior to tissue collection. Sham laparotomy was used as a negative control. Group sizes: Sham WT, N = 6; Sham KO, N = 6, IRI WT, N = 16; and IRI KO, N = 8. Liver neutrophils and macrophages were respectively quantified by counting the number of (**A**) Ly6G-positive and (**B**) F4/80-positive cells per square millimeter of liver tissue. Data points represent individual animals. *P*-values were determined by ANOVA with Šidák’s multiple comparion.

**Supplemental Figure 4.**
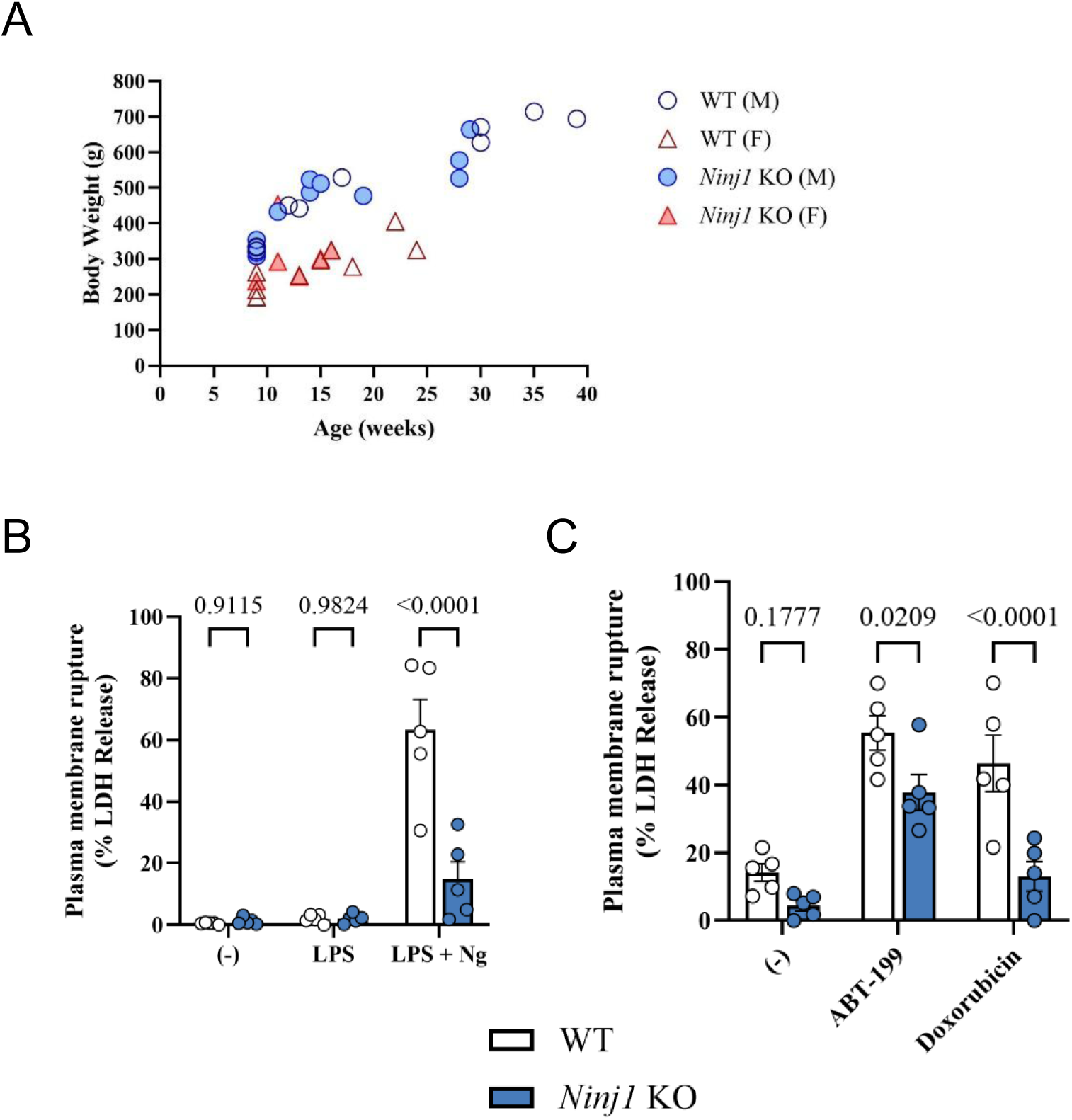
Baseline characteristics of Ninj1 knockout rats as compared to wild-type Sprague Dawley rats. (**A**) Untreated wild-type and *Ninj1* knockout male and Female rats weighed over time. Each datapoint represents an individual rat. (**B**) Bone marrow-derived macrophages were harvested from wild-type and *Ninj1* knockout rats. Pyroptosis was induced using lipopolysaccharide (100 ng/mL for 3 hours) and nigericin (20 µM for 30 min). (**C**) Secondary necrosis was induced using either ABT-199 (25 µM for 16 hours) or doxorubicin (10 µM for 16 hours). LDH release into the supernatant was measured to evaluate the extent of plasma membrane rupture. Data points represent BMDM harvested from different animals. *P*-values were determined by ANOVA.

**Supplemental Figure 5.**
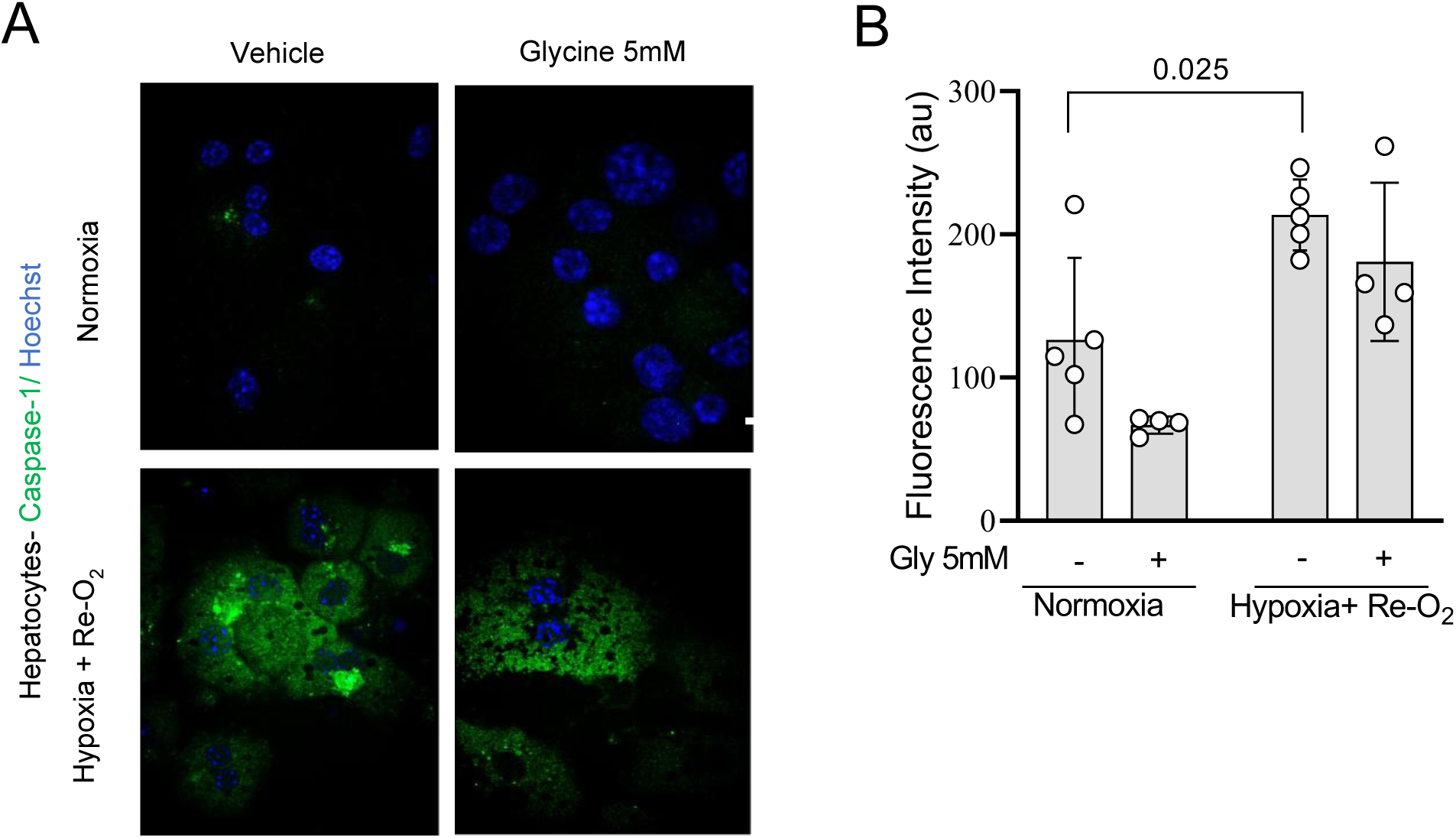
NINJ1 inhibition does not affect upstream caspase-1 activation in primary mouse hepatocytes subjected to hypoxia-reoxygenation. (**A**) Representative confocal images of primary hepatocytes labeled with FLICA (FAM-YVAD-FMK) to detect active caspase-1. Under normoxic conditions (21% O_2_), vehicle-treated cells and cells co-treated with glycine (5 mM) showed minimal FLICA signal. Following hypoxia–reoxygenation, hepatocytes exhibited marked caspase-1 activation, which remained detectable in both vehicle and glycine-treated groups, indicating that NINJ1 inhibition does not alter upstream caspase-1 activation. (**B**) Quantification of FLICA fluorescence intensity confirms increased caspase-1 activation under hypoxia–reoxygenation in both vehicle- and glycine-treated hepatocytes, with no reduction in response to NINJ1 inhibition.

**Supplemental Table 1.** Liver transplant patient demographic and clinical data.

| Patient number | EAD (Y/N) | Sex | MELD | Primary diagnosis | Secondary diagnosis | Donor age | Transplant warm ischemic time | Transplant cold ischemic time | AST |
| --- | --- | --- | --- | --- | --- | --- | --- | --- | --- |
| 1 | Y | Male | 14 | Alcoholic Cirrhosis | -- | [unavailable] | 72 | 788 | 32167 |
| 2 | Y | Male | 15 | Alcoholic Cirrhosis | HPS | 46 | 43 | 296 | 9408 |
| 3 | Y | Male | 26 | Alcoholic Cirrhosis | -- | 36 | 42 | 272 | 7891 |
| 4 | Y | Male | 19 | Retransplant | HCC | 58 | 12 | 672 | 6951 |
| 5 | Y | Male | 7 | Hepatitis C | HCC | [unavailable] | 14 | 599 | 6603 |
| 6 | Y | Male | 8 | Alcoholic Cirrhosis | HCC | 53 | 16 | 335 | 6395 |
| 7 | Y | Female | 7 | Hepatitis C | HCC | 46 | 14 | 211 | 6323 |
| 8 | Y | Male | 29 | Alcoholic Cirrhosis | -- | 45 | 57 | 443 | 7864 |
| 9 | Y | Male | 17 | Hepatitis C | HCC | 48 | 43 | 326 | 8654 |
| 10 | Y | Male | 16 | Alcoholic Cirrhosis | -- | 23 | 43 | 234 | 25889 |
| 11 | Y | Male | 9 | Alcoholic Cirrhosis | HCC | 16 | 78 | 398 | 5197 |
| 12 | N | Female | [unavailable] | α1 - Antitrypsin Deficiency | -- | 20 | 29 | 312 | 102 |
| 13 | N | Male | 6 | Hepatitis B | HCC | 21 | 27 | 301 | 481 |
| 14 | N | Male | [unavailable] | Cirrhosis | MASH | 65 | 13 | 312 | 162 |
| 15 | N | Male | 14 | Hepatitis C | Alcoholic Cirrhosis | 41 | 52 | 530 | 170 |
| 16 | N | Male | 34 | Alcoholic Cirrhosis |  | 69 | 45 | 273 | 114 |
| 17 | N | Male | 32 | Cirrhosis | MASH | 31 | 45 | 315 | 286 |
| 18 | N | Female | 13 | Retransplant | PSC | 18 | 48 | 312 | 138 |

Abbreviations: Aspartate transaminase, AST; Early allograft dysfunction, EAD; Hepatocellular carcinoma, HCC; Hepatopulmonary syndrome, HPS; Metabolic dysfunction-associated steatohepatitis, MASH. Early allograft dysfunction was defined as an AST value >5000 IU/mL, consistent with standard definitions (28, 64).

## References

1. Galluzzi L, et al. Molecular mechanisms of cell death: recommendations of the Nomenclature Committee on Cell Death 2018. Cell Death Differ. 2018;25(3):486–541.

2. Newton K, et al. Cell death. Cell. 2024;187(2):235–256.

3. Hirao H, Nakamura K, Kupiec-Weglinski JW. Liver ischaemia–reperfusion injury: a new understanding of the role of innate immunity. Nat Rev Gastroenterol Hepatol. 2022;19(4):239–256.

4. Liu J, Man K. Mechanistic Insight and Clinical Implications of Ischemia/Reperfusion Injury Post Liver Transplantation. Cell Mol Gastroenterol Hepatol. 2023;15(6):1463–1474.

5. Hartleif S, et al. Long-term Outcome of Asymptomatic Patients With Graft Fibrosis in Protocol Biopsies After Pediatric Liver Transplantation. Transplantation. 2023;107(11):2394–2405.

6. Gautheron J, Gores GJ, Rodrigues CMP. Lytic cell death in metabolic liver disease. J Hepatol. 2020;73(2):394–408.

7. Kondo T, et al. The role of RIPK1 mediated cell death in acute on chronic liver failure. Cell Death Dis. 2021;13(1):5.

8. Khanova E, et al. Pyroptosis by caspase11/4-gasdermin-D pathway in alcoholic hepatitis in mice and patients. Hepatology. 2018;67(5):1737–1753.

9. Kolachala VL, et al. Ischemia reperfusion injury induces pyroptosis and mediates injury in steatotic liver thorough Caspase 1 activation. Apoptosis. 2021;26(5–6):361–370.

10. Li J, et al. Blocking GSDMD processing in innate immune cells but not in hepatocytes protects hepatic ischemia-reperfusion injury. Cell Death Dis. 2020;11(4):244.

11. Guicciardi ME, et al. Apoptosis and necrosis in the liver. Compr Physiol. 2013;3(2):977–1010.

12. Broz P, Pelegrín P, Shao F. The gasdermins, a protein family executing cell death and inflammation. Nat Rev Immunol. 2020;20(3):143–157.

13. Zhong W, et al. Aging aggravated liver ischemia and reperfusion injury by promoting hepatocyte necroptosis in an endoplasmic reticulum stress-dependent manner. Ann Transl Med. 2020;8(14):869.

14. Wu J, et al. Ferroptosis in liver disease: new insights into disease mechanisms. Cell Death Discov. 2021;7(1):276.

15. Wu K, et al. STAT1 promotes ferroptosis and inflammation in mouse hepatic ischemia-reperfusion injury. Commun Biol. 2025;8(1):1549.

16. Lin J, et al. ZBP1-sensed hypoxic stress triggers intrinsic necroptosis in hepatocytes, aggravating hepatic ischemia-reperfusion injury: an experimental study. Int J Surg. 2025;111(11):7761–7776.

17. Kayagaki N, et al. NINJ1 mediates plasma membrane rupture during lytic cell death. Nature. 2021;591(7848):131–136.

18. Ramos S, et al. NINJ1 induces plasma membrane rupture and release of damage-associated molecular pattern molecules during ferroptosis. EMBO J. 2024;43(7):1164–1186.

19. Dondelinger Y, et al. NINJ1 is activated by cell swelling to regulate plasma membrane permeabilization during regulated necrosis. Cell Death Dis. 2023;14(11):755.

20. Araki T, Milbrandt J. Ninjurin, a novel adhesion molecule, is induced by nerve injury and promotes axonal growth. Neuron. 1996;17(2):353–361.

21. Jennewein C, et al. Contribution of Ninjurin1 to Toll-like receptor 4 signaling and systemic inflammation. Am J Respir Cell Mol Biol. 2015;53(5):656–663.

22. Degen M, et al. Structural basis of NINJ1-mediated plasma membrane rupture in cell death. Nature. 2023;618(7967):1065–1071.

23. Sahoo B, et al. How NINJ1 mediates plasma membrane rupture and why NINJ2 cannot. Cell. 2025;188(2):292–302.e11.

24. David L, et al. NINJ1 mediates plasma membrane rupture by cutting and releasing membrane disks. Cell. 2024;187(9):2224–2235.e16.

25. Kayagaki N, et al. Inhibiting membrane rupture with NINJ1 antibodies limits tissue injury. Nature. 2023;618(7967):1072–1077.

26. Borges JP, et al. Glycine inhibits NINJ1 membrane clustering to suppress plasma membrane rupture in cell death. eLife. 2022;11:e78609.

27. den Hartigh AB, et al. Muscimol inhibits plasma membrane rupture and ninjurin-1 oligomerization during pyroptosis. Commun Biol. 2023;6(1):1010.

28. Wiemann BA, et al. Early Allograft Dysfunction after liver transplantation- definition, incidence and relevance in a single-centre analysis. Langenbecks Arch Surg. 2025;410(1):76.

29. Yue S, et al. Prolonged Ischemia Triggers Necrotic Depletion of Tissue-Resident Macrophages To Facilitate Inflammatory Immune Activation in Liver Ischemia Reperfusion Injury. The Journal of Immunology. 2017;198(9):3588–3595.

30. Pourmal S, et al. Autoinhibition of dimeric NINJ1 prevents plasma membrane rupture. Nature. 2024;637(8045):446–452.

31. Suzuki S, et al. Neutrophil infiltration as an important factor in liver ischemia and reperfusion injury. Modulating effects of FK506 and cyclosporine. Transplantation. 1993;55(6):1265–1272.

32. Datta G. Molecular mechanisms of liver ischemia reperfusion injury: Insights from transgenic knockout models. WJG. 2013;19(11):1683.

33. Fink SL, Cookson BT. Apoptosis, pyroptosis, and necrosis: mechanistic description of dead and dying eukaryotic cells. Infect Immun. 2005;73(4):1907–1916.

34. Bachmann S, et al. Glycine in Carolina rinse solution reduces reperfusion injury, improves graft function, and increases graft survival after rat liver transplantation. Transplant Proc. 1995;27(1):741–742.

35. Duenschede F, et al. Different protection mechanisms after pretreatment with glycine or alpha-lipoic acid in a rat model of warm hepatic ischemia. Eur Surg Res. 2006;38(6):503–512.

36. Gassner JMGV, et al. Improvement of Normothermic Ex Vivo Machine Perfusion of Rat Liver Grafts by Dialysis and Kupffer Cell Inhibition With Glycine. Liver Transpl. 2019;25(2):275–287.

37. Sheth H, et al. Glycine maintains mitochondrial activity and bile composition following warm liver ischemia-reperfusion injury. J Gastroenterol Hepatol. 2011;26(1):194–200.

38. Yamanouchi K, et al. Glycine reduces hepatic warm ischaemia-reperfusion injury by suppressing inflammatory reactions in rats. Liver Int. 2007;27(9):1249–1254.

39. Zhong Z, Jones S, Thurman RG. Glycine minimizes reperfusion injury in a low-flow, reflow liver perfusion model in the rat. Am J Physiol. 1996;270(2 Pt 1):G332–338.

40. Loomis WP, et al. Diverse small molecules prevent macrophage lysis during pyroptosis. Cell Death Dis. 2019;10(4):326.

41. Baratta JL, et al. Cellular organization of normal mouse liver: a histological, quantitative immunocytochemical, and fine structural analysis. Histochem Cell Biol. 2009;131(6):713–726.

42. Ellett JD, et al. Murine Kupffer cells are protective in total hepatic ischemia/reperfusion injury with bowel congestion through IL-10. J Immunol. 2010;184(10):5849–5858.

43. Zhang L, et al. Activation of NLRP3 Inflammasome via Drp1 Overexpression in Kupffer Cells Aggravates Ischemia-reperfusion Injury in Hepatic Steatosis. J Clin Transl Hepatol. 2023;000(000):000–000.

44. Evans ZP, et al. Mitochondrial uncoupling protein-2 deficiency protects steatotic mouse hepatocytes from hypoxia/reoxygenation. Am J Physiol Gastrointest Liver Physiol. 2012;302(3):G336–342.

45. Hu Y, et al. The Ninj1/Dusp1 Axis Contributes to Liver Ischemia Reperfusion Injury by Regulating Macrophage Activation and Neutrophil Infiltration. Cell Mol Gastroenterol Hepatol. 2023;15(5):1071–1084.

46. Culemann S, et al. Stunning of neutrophils accounts for the anti-inflammatory effects of clodronate liposomes. J Exp Med. 2023;220(6):e20220525.

47. Culemann S, et al. Addendum: Stunning of neutrophils accounts for the anti-inflammatory effects of clodronate liposomes. J Exp Med. 2023;220(7):e2022052506022023a.

48. Volchuk A, et al. Indirect regulation of HMGB1 release by gasdermin D. Nat Commun. 2020;11(1):4561.

49. Schachter J, et al. Gasdermin D mediates a fast transient release of ATP after NLRP3 inflammasome activation before ninjurin 1-induced lytic cell death. Cell Rep. 2025;44(2):115233.

50. Tsung A, et al. The nuclear factor HMGB1 mediates hepatic injury after murine liver ischemia-reperfusion. J Exp Med. 2005;201(7):1135–1143.

51. Tanaka K, et al. Recipient toll-like receptor 4 determines the outcome of ischemia-reperfusion injury in steatotic liver transplantation in mice. Am J Transplant. 2025;25(6):1168–1179.

52. Sosa RA, et al. Disulfide High-Mobility Group Box 1 Drives Ischemia-Reperfusion Injury in Human Liver Transplantation. Hepatology. 2021;73(3):1158–1175.

53. Terry AQ, et al. Disulfide-HMGB1 signals through TLR4 and TLR9 to induce inflammatory macrophages capable of innate-adaptive crosstalk in human liver transplantation. Am J Transplant. 2023;23(12):1858–1871.

54. Zhang Q, et al. Circulating mitochondrial DAMPs cause inflammatory responses to injury. Nature. 2010;464(7285):104–107.

55. Westhaver LP, et al. Mitochondrial DNA levels in perfusate and bile during ex vivo normothermic machine correspond with donor liver quality. Heliyon. 2024;10(5):e27122.

56. Yoshino O, et al. Elevated levels of circulating mitochondrial DNA predict early allograft dysfunction in patients following liver transplantation. J Gastroenterol Hepatol. 2021;36(12):3500–3507.

57. Pollara J, et al. Circulating mitochondria in deceased organ donors are associated with immune activation and early allograft dysfunction. JCI Insight. 2018;3(15):e121622, 121622.

58. Zhu Y, et al. NINJ1 regulates plasma membrane fragility under mechanical strain. Nature. 2025;644(8078):1088–1096.

59. Araki T, et al. Mechanism of homophilic binding mediated by ninjurin, a novel widely expressed adhesion molecule. J Biol Chem. 1997;272(34):21373–21380.

60. Dutkowski P, et al. Challenges to liver transplantation and strategies to improve outcomes. Gastroenterology. 2015;148(2):307–323.

61. Ito T, et al. Ischemia-reperfusion injury and its relationship with early allograft dysfunction in liver transplant patients. Am J Transplant. 2021;21(2):614–625.

62. Czigany Z, et al. Machine perfusion for liver transplantation in the era of marginal organs-New kids on the block. Liver Int. 2019;39(2):228–249.

63. Goldaracena N, et al. Expanding the donor pool for liver transplantation with marginal donors. Int J Surg. 2020;82S:30–35.

64. Agopian VG, et al. Evaluation of Early Allograft Function Using the Liver Graft Assessment Following Transplantation Risk Score Model. JAMA Surg. 2018;153(5):436–444.

65. Andrews TS, et al. Single-cell, single-nucleus, and spatial transcriptomics characterization of the immunological landscape in the healthy and PSC human liver. J Hepatol. 2024;80(5):730–743.

66. Selzner N, et al. Ischemic preconditioning protects the steatotic mouse liver against reperfusion injury: an ATP dependent mechanism. J Hepatol. 2003;39(1):55–61.

67. Percie Du Sert N, et al. Reporting animal research: Explanation and elaboration for the ARRIVE guidelines 2.0. PLoS Biol. 2020;18(7):e3000411.

68. Charni-Natan M, Goldstein I. Protocol for Primary Mouse Hepatocyte Isolation. STAR Protocols. 2020;1(2):100086.

69. Wittig I, Braun H-P, Schägger H. Blue native PAGE. Nat Protoc. 2006;1(1):418–428.

